# New SAR11 isolate genomes and global marine metagenomes resolve ecologically relevant units within the *Pelagibacterales*

**DOI:** 10.1101/2024.12.24.630191

**Authors:** Kelle C. Freel, Sarah J. Tucker, Evan B. Freel, Ulrich Stingl, Stephen J. Giovannoni, A. Murat Eren, Michael S. Rappé

## Abstract

The bacterial order *Pelagibacterales* (SAR11) is widely distributed across the global surface ocean, where its activities are integral to the marine carbon cycle. High-quality genomes from isolates that can be propagated and phenotyped are needed to unify perspectives on the ecology and evolution of this complex group. Here, we increase the number of complete SAR11 isolate genomes threefold by describing 81 new SAR11 strains from coastal and offshore surface seawater of the tropical Pacific Ocean. Our analyses of the genomes and their spatiotemporal distributions support the existence of 29 monophyletic, discrete *Pelagibacterales* ecotypes that we define as genera. The spatiotemporal distributions of genomes within genera were correlated at fine scales with variation in ecologically-relevant gene content, supporting generic assignments and providing indications of speciation. We provide a hierarchical system of classification for SAR11 populations that is meaningfully correlated with evolution and ecology, providing a valid and utilitarian systematic nomenclature for this clade.

## Introduction

SAR11 marine bacteria, known as the *Pelagibacterales*^1^, are a genetically diverse, order-level lineage of heterotrophs within the *Alphaproteobacteria.* They numerically dominate planktonic communities across the global ocean^2–6^. Correlations between the spatiotemporal distributions of SAR11 lineages, environmental variables, and the subclade structure of *Pelagibacterales*, have long indicated the presence of distinct ecotypes within SAR11^3,7–10^. By linking functional traits with phylogenetic and biogeographic patterns, our recent findings support the differentiation of SAR11 into functionally distinct sublineages that segregate over short spatial scales^11^. High-quality SAR11 genomes, available early in the investigation of SAR11 diversity, demonstrated they are a remarkably cohesive genetic assemblage^1^, but were too few to discern complex patterns in the adaptive machinery underlying the stunning success of these minimalist microorganisms.

Since the first observation of SAR11 through environmental 16S rRNA gene fragments over three decades ago^12^, microbiology has benefited from a dramatic increase in microbial sequence data recovered directly from the environment, offering representative genomes for many difficult to cultivate microbial lineages^13^. However, even extensive genome-resolved surveys of marine metagenomes have failed to yield high-quality SAR11 genomes^14^, resulting in limited insights into what constitutes ecologically meaningful units within this broad group. The extensive intra-clade diversity of SAR11^15–17^ confounds the ability to reconstruct environmental genomes from metagenomes^18,19^, which is why one of the most abundant microbial clades in marine systems suffers from poor representation in genome-resolved metagenomics surveys^20^. By circumventing the need to assemble complex metagenomes to recover genomes, single-cell sorting coupled with whole-genome amplification techniques have been much more effective in sampling environmental SAR11 populations. However, in an extensive effort to characterize surface ocean microbes, the estimated genome completion of single-amplified genomes (SAGs) that could be affiliated with SAR11 remained below 60%^21^, a level that prevents robust phylogenomic insights. Such barriers have led to a reliance on isolate genomes to investigate the evolution of SAR11 populations^1,22–25^, yet following this path has been impeded by another formidable challenge: the difficulty of cultivating SAR11 in the laboratory, even with genomic insights regarding its unique growth requirements^26–28^.

The first successful cultivation of SAR11 in 2002 resulted in the isolation of *Pelagibacter* strain HTCC1062^29^, followed by the publication of its complete genome^30^. Over the past two decades, additional isolate genomes have been few, with only 28 currently available. Despite their rarity, high-quality genomes from isolated strains not only shed light on SAR11 biology^26,31,32^ and the origins of this lineage within the *Alphaproteobacteria*^1,22,23^, but also have made it possible to establish key concepts in biology such as genome streamlining^1,31–36^ and investigate the evolutionary processes that shape protein evolution^8,16^.

Here we report 81 high-quality genomes from SAR11 strains, increasing the number available for SAR11 isolates by threefold, and leverage this new collection to build a robust genome phylogeny for the order *Pelagibacterales*. By incorporating publicly available, high-quality single-cell genomes and surface ocean metagenomes from both a steep, nearshore to open-ocean local environmental gradient and elsewhere from around the globe, we reveal cohesive patterns of genomic and ecotypic diversification. We propose a framework through which to characterize and interpret genome heterogeneity at multiple stages along the evolutionary history of SAR11 marine bacteria, and establish a roadmap for future efforts to organize this globally abundant bacterial clade.

## Results

### Eighty-one high-quality genomes sequenced from 206 newly isolated SAR11 strains and co-cultures

Three dilution-to-extinction culturing experiments using surface seawater collected from nearshore and adjacent offshore environments of Oʻahu, Hawaiʻi, in the tropical Pacific Ocean resulted in 916 isolates from 2,102 inoculated cultures (Table 1; Supplementary Fig. 1). Using a streamlined isolate-to-genome approach, we identified 206 cultures as either pure SAR11 strains or mixed cultures with at least 50% of the total reads matching a SAR11 strain via 16S rRNA gene amplicon sequencing (Supplementary Data 1), and sequenced draft genomes from 90. Manual curation resulted in 79 high-quality SAR11 isolate genomes. The genomes from two strains (HIMB123 and HIMB109) isolated from a previous culture experiment were also added^37^, resulting in 81 new SAR11 genomes from isolates. The majority of these (n=60) assembled into ten contigs or less, including 24 closed genomes with an additional 30 containing one to three contigs. They ranged from 1.00 to 1.54 Mbp in size with a GC content of 28.5 to 30.7% (Supplementary Data 2). The median pairwise genome-wide average nucleotide identity (gANI) value across all genomes was 81.8% and none of the 81 new isolate genomes were identical. Having captured a genetically diverse array of SAR11 isolates, we used a phylogenomic approach to characterize evolutionary relationships among these genomes and high-quality single-cell and isolate genomes previously retrieved from seawater.

**Table 1.** Summary of high-throughput culturing (HTC) experiments.

| Site | Inoculum source | Inoculum size<br>(# of cells) | Cultures<br>screened | Positive<br>cultures | SAR11<br>genomes |
| --- | --- | --- | --- | --- | --- |
| SB | raw seawater | 5 | 576 | 339 | 53 |
| STO1 | raw seawater | 5 | 576 | 126 | 16 |
| STO1 | cryopreserved seawater | 5 | 480 | 142 | 9 |
| STO1 | cryopreserved seawater | 100 | 470 | 343 | 1 |

### An extensive genome phylogeny reveals a robust evolutionary backbone populated by clusters of closely related genomes

We first sought to resolve relationships between the strains isolated in this study and other publicly available high-quality *Pelagibacterales* genomes to precisely establish where the new genomes originate from within the broad spectrum of known SAR11 diversity. For this, we created a database that, in addition to the 81 genomes presented here, included 28 public SAR11 isolate genomes, 8 of which were also isolated from off the windward coast of Oʻahu, Hawaiʻi, and 375 SAR11 single-amplified genomes (SAGs) estimated to be ≥85% complete with a redundancy <5% (Supplementary Data 3; Supplementary Notes). We also included five additional SAR11 SAGs of potentially unique evolutionary origin within the collection of 375^21,38,39^. This resulted in a curated collection of 484 SAR11 genomes to assess the evolutionary backbone for SAR11.

Previous studies investigating phylogenomic relationships within the *Alphaproteobacteria* utilized a curated set of 200 single-copy core genes (SCGs) for this bacterial class^23,40^. We evaluated the presence of these 200 SCGs across our genome dataset, and excluded genes missing in more than 90% of the 484 SAR11 genomes or redundant in more than 2% of the genomes. This resulted in a SAR11-specific SCG set of 165 genes that were translated to amino acid sequences for downstream phylogenomic analyses, referred to hereafter as the SAR11_165 protein sequence set (Supplementary Data 4).

Our analysis of the 484 genomes using the SAR11_165 protein sequence set revealed that the SAR11 clade consists of four robust, deeply-branching sublineages (Fig. 1; Supplementary Fig. 2). Three of these branches were the previously characterized subclades Ic^38^, II^41^, and III^1,38^, while the fourth was a combination of established SAR11 subclades Ia and Ib^41^, which did not form separate monophyletic subclades in this exhaustive genomic dataset and phylogenetic analysis. If the SAR11 clade is assigned to the taxonomic level of a bacterial order^1^, then these four lineages logically resolve to the taxonomic level of families. There is compelling evolutionary and comparative genomic evidence that the AEGEAN-169 lineage, also known previously as SAR11 subclade V^1,22,42^ does not share common ancestry with other SAR11 genomes^23,43,44^. While less studied than AEGEAN-169, putative SAR11 subgroup IV^38^ is also unlikely to share common ancestry with SAR11^10,22,43,45^. Phylogenomic analyses that included genomes from these two groups reinforced their distant relationship to the core set of sublineages recognized as SAR11 (Supplementary Fig. 3).

**Figure 1.**
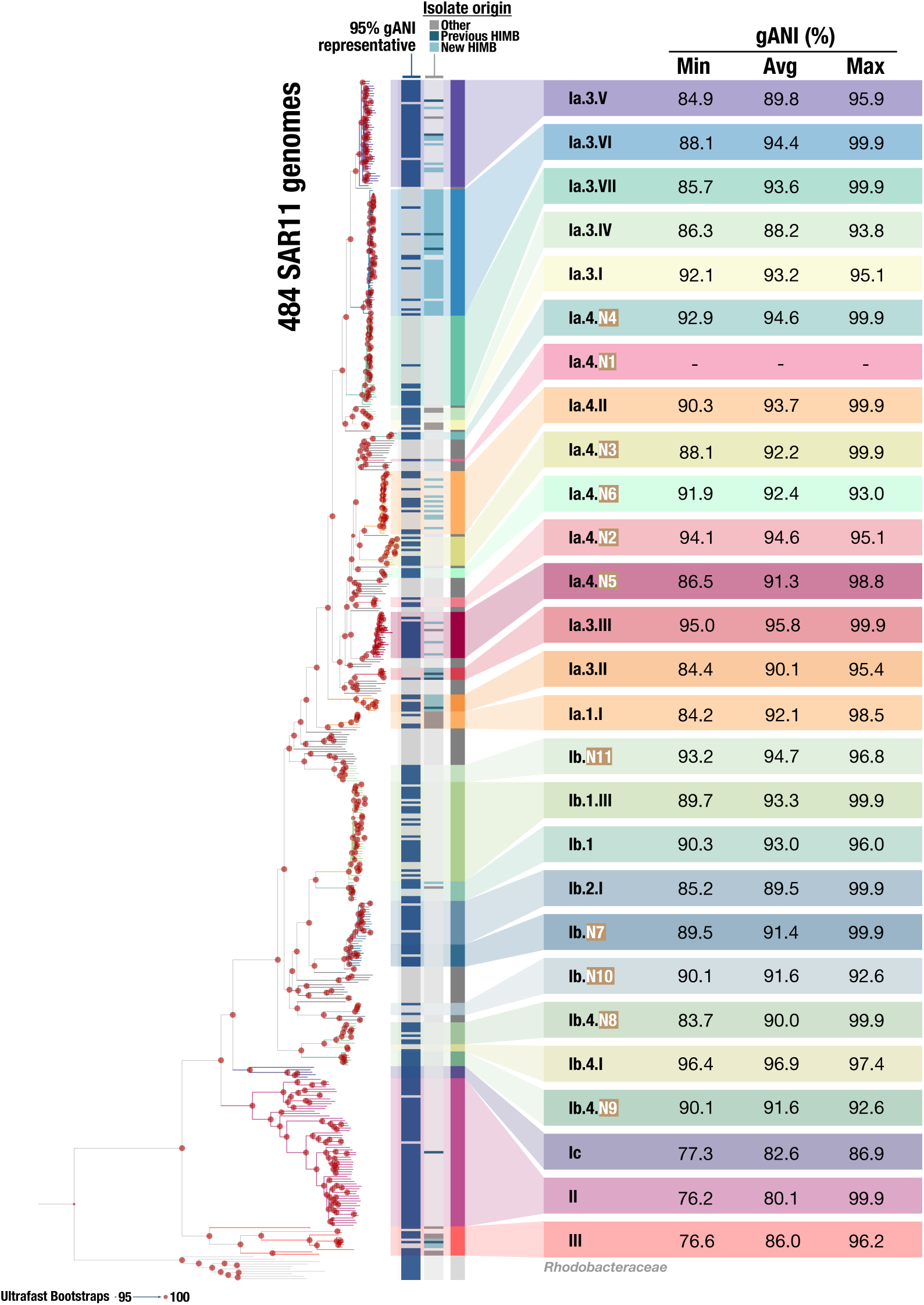
Phylogeny of the *Pelagibacterales*. A phylogeny of 484 SAR11 genomes (109 isolates and 375 SAGs) based on a concatenated alignment of protein sequences from 165 SAR11 core genes. Also included are summaries of the minimum, average, and maximum gANI for each clade. Genomes identified as clade representatives using a 95% ANI threshold are highlighted with a dark blue bar in the indicated column. The isolate origin indicates new isolate genomes published as part of this study originating from Kāneʻohe Bay and cultivated at the Hawaiʻi Institute of Marine Biology (HIMB) (light blue), previous isolate genomes from the HIMB collection (dark blue), or isolate genomes published from other sources (gray).

We further removed genomes from this initial tree in two steps. First we excluded SAGs that did not fall into a 90% gANI cluster of at least three genomes to focus our analyses on well-resolved regions of the tree. Second, we de-replicated the remaining genomes using a conservative cutoff of 95% gANI (Supplementary Data 5) to minimize subsequent competitive metagenomic recruitment steps splitting reads among closely related genomes^46^. While the 95% ANI cutoff is broadly recognized in contemporary microbiology as a threshold to identify microbial species, it over-splits ecologically and evolutionarily cohesive units in SAR11 and does not delineate species-like groups^8,15,17,47^. We note that the reason behind our use of the 95% gANI in this step of our analysis was solely to establish a technically robust workflow prior to competitive read recruitment rather than a biologically meaningful partitioning of our genomes, a challenge our study focuses on later.

We then turned our attention to the distal end of the phylogeny, which contained a large number of well-supported clusters of closely related genomes, particularly within the Ia/Ib subgroup that contained 78 of the 81 new isolate genomes. Focusing on well-supported clades within the phylogeny revealed 24 monophyletic clusters within the historical Ia/Ib subgroup that were characterized by a range of gANI values from 84% to 96% (92.1 ± 2.9%; mean ± SD) (Fig. 1). While some of these clusters were recognized previously, we defined an additional 11 here (Fig 1; Supplementary Data 6). Twelve of the 24 clusters contained an isolated representative, and eight contained at least one isolate from our study area in the tropical Pacific.

In summary, our extensive phylogenomic analysis of SAR11 revealed 24 monophyletic clusters within the historical Ia/Ib subgroup which included the majority of SAR11 SAGs and the new and previously published isolate genomes. The non-uniform minimum gANI estimates suggest that the application of sequence-based ANI thresholds to demarcate SAR11 diversity may obscure important evolutionary signals. Hypothesized drivers of the maintenance and partitioning of genomic diversity in SAR11 include niche-based processes, where genetically cohesive clusters also display ecological homogeneity and the underlying genetic diversity is maintained by similar forces of selection, recombination, and drift^24,34,48^. To understand the potential eco-evolutionary forces that shape SAR11 diversification, we turned to metagenomic read recruitment analysis to recover biogeographical distribution patterns for our genomes across the globe.

### Global read recruitment from the surface ocean reveals broadly congruent phylogenetic and ecotypic diversification across SAR11

Our competitive metagenomic read recruitment assessed the distribution of 268 SAR11 genomes (dereplicated at 95% gANI from the initial set of 484) around the globe and relied upon 1,345 publicly-available marine metagenomes, as well as metagenomes from Kāneʻohe Bay via the Kāneʻohe Bay Time-series (KByT)^9^, the location of isolation for the 81 new and 9 of the 28 existing isolate genomes (Supplementary Data 7; Supplementary Data 8). SAR11 genomes collectively recruited a total of 5,074,677,192 reads from the entire collection of global metagenomes that contained a total of 70,533,279,194 reads, or 7.2% of the dataset. The SAR11 genomes accounted for 0.001% to 27.7% of reads in each metagenome with a median genome coverage up to 12x (Supplementary Data 8, Supplementary Data 9). These data enabled us to investigate whether cohesive genomic and ecological groups, or ecotypes, could be discerned by combining SAR11 phylogeny and biogeography.

Our first priority was to establish whether genome clusters within a given SAR11 clade showed cohesive read recruitment profiles across metagenomes, or, in other words, whether the ecological patterns revealed by a single genome were similar to all genomes within the group to which it belonged. We narrowed our focus here to the historical subgroups Ia/Ib where the majority of isolated cultures are classified. Detection values (proportion of nucleotides in a genome that are covered at least 1x) for multiple genomes within a genome cluster showed a significant degree of cohesion (PERMANOVA, p < 0.001, Supplementary Fig. 4; Fig. 2; Supplementary Data 10). For example, patterns of detection for representatives from Ia.3.IV, Ia.3.I, Ia.4.II, Ia.4.N2, Ia.4.N5, Ib.1.III, and Ib.4.N9 show particularly consistent within-cluster patterns (Fig. 2, Supplementary Fig. 4). Overlap in patterns of detection between SAGs and isolate genomes within the same clade demonstrated that both genome types accurately reflect distribution patterns for closely related populations as inferred by phylogeny (Fig. 2, Supplementary Data 12). A non-metric multidimensional scaling (NMDS) analysis of the overall detection patterns of genomes across metagenomes consistently grouped genomes within a given clade more closely compared to those that belonged to other genome clusters (Supplementary Fig. 4), further supporting a high degree of intra-clade ecological cohesion.

**Figure 2:**
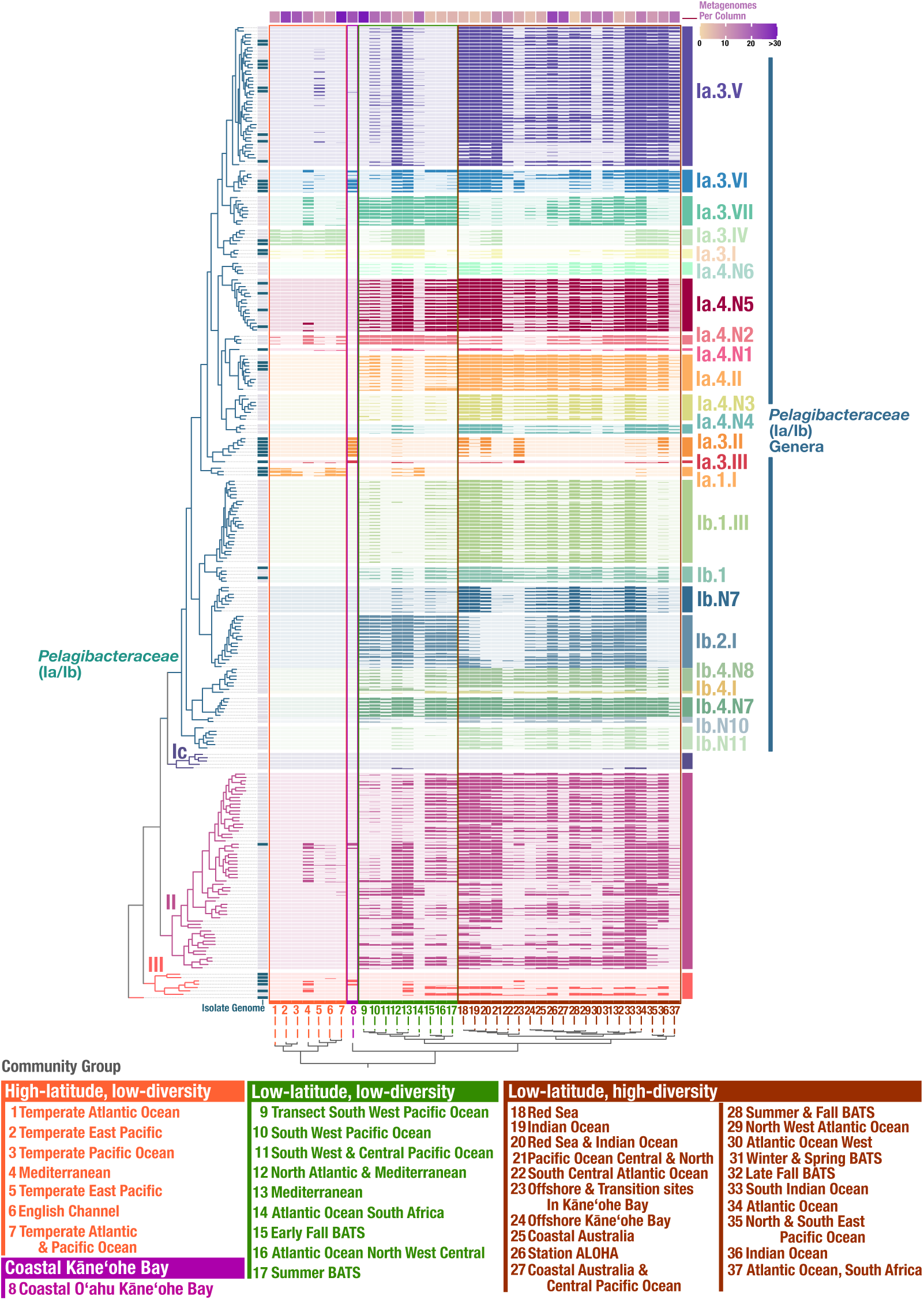
Global metagenome read recruitment to 268 *Pelagibacterales* genomes. A clustering analysis reveals that the distribution of metagenomes from the same geographic location have characteristic patterns of detection. Each row represents the average detection values for a given genome in the metagnome cluster, with the number of metagenomes averaged per column indicated at the top. A total of 552 metagnomes are included (Supplemental Table 11). The community groups are defined and color coded as: “High-latitude, low-diversity” (orange), “Coastal Kāneʻohe Bay” (pink), “Low-lattitude, low-diversity” (green), and “Low-lattitude, high diversity” (red). Detection values from 0.25 to 0.75 are shown.

Our second priority was to establish insights into whether SAR11 genome clusters differed in their biogeographical patterns, and whether genome clusters identified SAR11 populations of distinct ecology. Hierarchical clustering of metagenomes based on detection patterns of the SAR11 community revealed four groups: metagenomes that originated from (1) low-latitude samples with low SAR11 diversity, (2) high-latitude samples with low SAR11 diversity, (3) low-latitude samples with high SAR11 diversity, and (4) samples from coastal Kāneʻohe Bay (Fig. 2). The distribution of SAR11 genomes differed significantly across these four community groups (PERMANOVA, p < 2.2e-16; Supplementary Fig. 5). Many SAR11 genome clusters were indeed differentially distributed across these community groups. For example, Ia.4.II and Ia.4.N5 were only consistently found in the low-latitude, low-diversity and low-latitude, high-diversity communities, while Ia.3.IV and Ia.I were found across the high-latitude, low-diversity communities and only in select sites in the other community groups (Fig. 2). However, in multiple cases, the environmental detection patterns of different phylogenomic genome clusters overlapped; while there was some degree of inter-clade ecological differentiation, distinct SAR11 genome clusters frequently co-occurred (Fig. 2, Supplementary Fig. 6, Supplementary Data 12). This observation suggests that patterns of distribution alone cannot discern the boundaries of cohesive, fundamental units within the ecotypically- and evolutionarily-diverse landscape of SAR11, a task that evidently requires the integration of biogeographical patterns identified using metagenomic read recruitment and ancestral relationships discerned by phylogenomics.

Finally, we used the read recruitment analysis to assign ecological patterns to specific SAR11 genome clusters. While multiple broad patterns were clear from the pairing of the phylogenomic relationships and read recruitment data, we focused our investigation on whether the genome clusters within the SAR11 Ia/Ib lineage that appeared to be confined to the coastal end of the KByT environmental gradient (Ia.3.VI, Ia.3.II, and Ia.3.III; Supplementary Fig. 1; also see^11^) were similarly constrained to coastal areas globally. Indeed, two of the three genome clusters, Ia.3.II and Ia.3.III, were detected almost exclusively in metagenomes sourced from coastal environments (e.g., Kāneʻohe Bay, the north coast of Panama, Chesapeake Bay, and the Atlantic coast of Portugal). Interestingly, while the clade Ia.3.VI was restricted to nearshore metagenomes across KByT, it was well-detected in both coastal and offshore environments in other oceanic regions (Fig. 2). Genome clusters Ia.3.II and Ia.3.III did not include any SAGs and were only composed of isolates from coastal Kāneʻohe Bay. Yet, we could detect them in rare instances in other oceans, which confirms their global relevance as representatives of SAR11 populations adapted to coastal ecosystems.

Through the combination of global metagenomic read recruitment and phylogenomics, we show that SAR11 genome clusters contain genomes with a high degree of intra-clade ecological cohesion. These genome clusters were often distinguished by their ecological distributions and demonstrated notable inter-clade ecological differentiation. Next, we applied this framework to understand how SAR11 genetic and ecological diversity partitions among ocean biomes, in particular coastal ocean and open ocean environments. Finally, we compared our delineation of these genome clusters to taxonomic structures proposed elsewhere (Supplementary Notes), and find that clustering of genomes presented in our study more closely reflects the ecological and evolutionary trajectories of this diverse lineage than the strict metric based approaches implemented by solely relying on relative evolutionary divergence (RED) values or the ranks assigned by GTDB.

The integrated ecological and evolutionary framework here is supported by high-quality genomes that span the known diversity of the *Pelagibacterales*, providing a critical opportunity to discern distinct ecologically meaningful genome clusters within SAR11. We show that the 24 distinct genome clusters within the combined subgroup Ia/Ib lineage represent groups sharing cohesive ecological patterns and evolutionary relationships, not at the finest tips of the phylogenomic tree, but at relatively deeper branches that encompass gANI values ranging from 84% to 96%. This level of genomic differentiation suggests it is plausible that these genome clusters represent SAR11 diversity at a higher taxonomic level than ‘species’. The cohesion of these genome clusters is further supported by recent work^11^, which reveals systematic differences in the metabolic potential of SAR11 genome clusters that likely support distinct ecological distributions in immediately adjacent coastal and open ocean surface seawater with habitat-specific metabolic genes that are under higher selective forces. With the combined evidence presented here and in the work by Tucker et al.^11^ that unite SAR11 diversity into distinct genome clusters with ecotype properties supported by SAR11 phylogenomics, ecology, metabolic potential, as well as population genetics, we argue that the most conceivable taxonomic rank at which SAR11 genome clusters can be described in a conventional framework emerges as the ‘genus’ level.

This genus-level designation is ideal as it encompasses a degree of diversity previously designated by SAR11 subgroups and has the flexibility to account for subtle variation in ecology recognized between closely related genomes. We identified the highest quality genome representatives (electing for isolates when possible) to assign as type genomes for each genus (Supplementary Data 13), which establishes a roadmap to rationally designate new genera as they are identified in the future.

### Evidence for ecological speciation within closely related genome clusters

Despite broad ecological cohesion within what we have designated as genera of the order *Pelagibacterales*, some notable differences highlight underlying complexities in defining the finest scales of divergence. The Ia.3.VI genus includes genomes from strains of Kāneʻohe Bay origin as well as SAGs from other regions of the global ocean, and encompasses significant genomic diversity (minimum gANI 88%) and phylogenomic structure (Fig. 3a). Through read recruitment (Fig. 2), we observed notable differences in the patterns of detection of Ia.3.VI genomes across community groups of (1) coastal Kāneʻohe Bay, (2) low-latitude, low-diversity, and (3) low-latitude, high-diversity. Examining these patterns more closely with a subset of metagenomes from Kāneʻohe Bay, BATS, and Station ALOHA (Fig. 3a,b), we found that isolate genomes from Kāneʻohe Bay harbored the highest detection values of the Ia.3.VI genus from metagenomes also sourced from the bay, while a SAG from the BATS site in the Atlantic Ocean (GCA_902533445.1) had the highest detection values at BATS (Fig. 3b), particularly in the summer and fall (Fig. 2).

**Figure 3:**
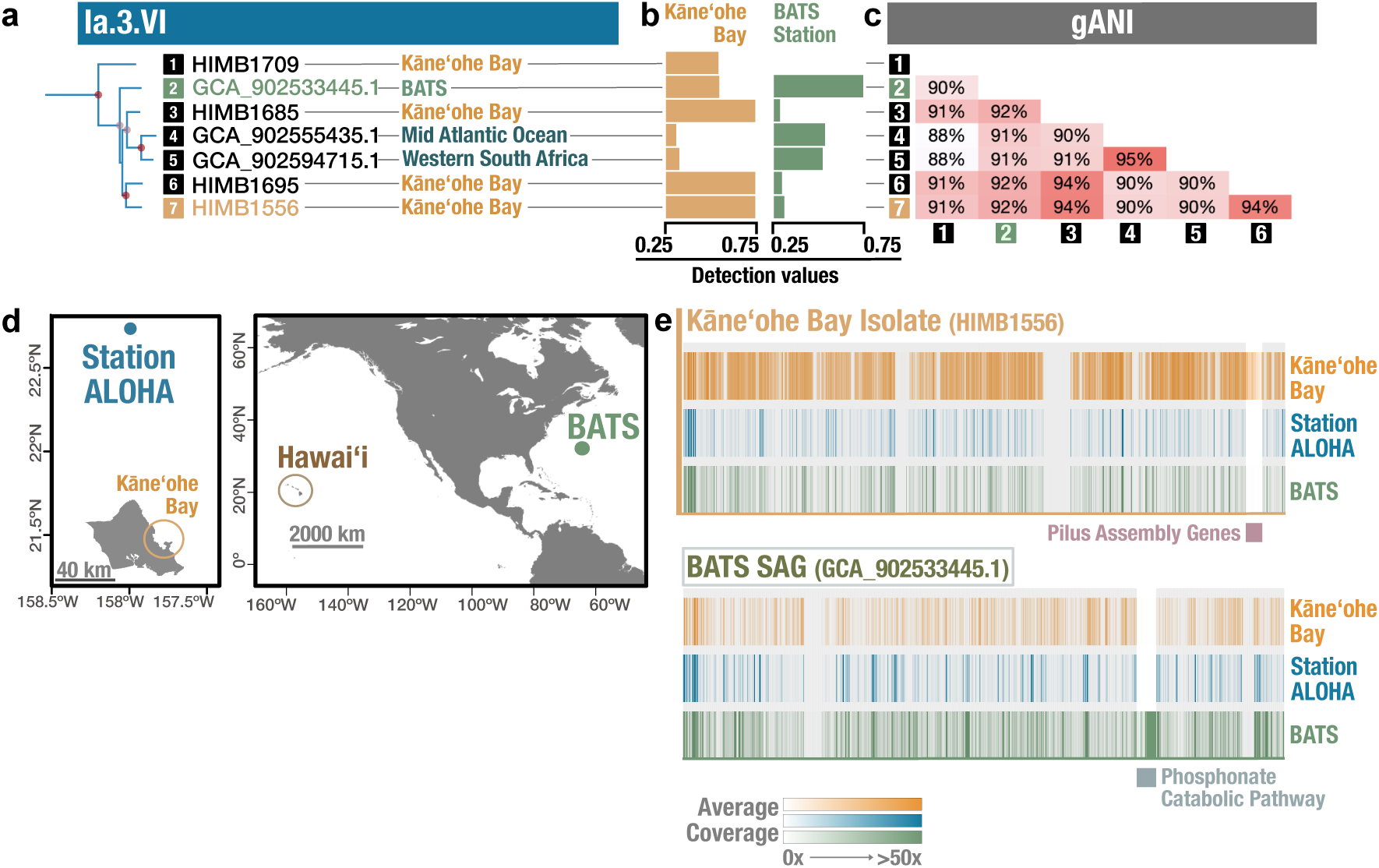
Fine-scale ecological speciation between closely related SAR11 genomes. (a) Subtree of Ia.3.VI genomes extracted from the full phylogeny shown in Figure 2, with geographic origins indicated. Circles at nodes indicate 98 (pink) and 100 (red) ultrafast bootstrap values. (b) Detection values, defined as the proportion of each genome with ≥1x coverage, are shown as averages across all nearshore Kāneʻohe Bay metagenomes (n=18) and surface (25 m) metagenomes from BATS (n=7). Values within the range of 0.25 and 0.75 are shown. (c) Matrix of pairwise whole genome average nucleotide identity (gANI) values. (d) Map of Oʻahu showing the locations of Kāneʻohe Bay and Station ALOHA (left), as well as the locations of the Hawaiian Islands and the Bermuda Atlantic Time-series (BATS) study site (right). (e) Coverage values across the genomes of isolate HIMB1556 and SAG GCA_902533445.1, averaged across metagenomes from nearshore Kāneʻohe Bay (n = 18), Station ALOHA in the North Pacific Subtropical Gyre (n = 12), and BATS in the Atlantic Ocean (n = 7), highlighting the differential detection of genes for type IV pilus assembly (HIMB1556) and phosphonate catabolic pathway (GCA_902533445.1).

Given the underlying genomic divergence between isolate HIMB1556 and BATS SAG GCA_902533445.1 and their distinct biogeographies, we next surveyed the genomes for unique gene content that may be relevant to their distributions. By inspecting the coverage of isolate HIMB1556 and BATS SAG GCA_902533445.1 using metagenomes from Kāneʻohe Bay and the BATS site, we found one genomic region of SAG GCA_902533445.1 that had particularly high coverage at BATS compared to the Kāneʻohe Bay samples (108.9x vs 0.2x ave. coverage; Supplementary Data 14). This region included 29 genes encoding the uptake (*phnCDE*) and catabolism of phosphonates via the C-P lyase pathway (*phnGHIJKLM*) (Fig. 3e; Supplementary Data 15) ^49^. The *phnJ* phylogeny did not reflect the phylogenomic relationships among genomes (Supplementary Fig. 7), and the pathway for phosphonate catabolism was located on a genomic island similar to a subset of genomes from Ia.3.I and Ia.3.VII^10,11,50^. The C-P lyase pathway is known to be enriched in phosphate-depleted systems of the Atlantic Ocean^51,52^ and Mediterranean Sea^10^, so the presence of C-P lyase catabolic genes in a genome sourced from BATS, but missing from a closely related genome sourced from the more phosphate-replete environment of Kāneʻohe Bay in the Pacific, suggests these genes provide an advantage in phosphate depleted systems and that the BATS SAG GCA_902533445.1 may be locally-adapted to its environment.

While the HIMB1556 genome lacked the C-P lyase pathway, it contained a unique genomic region with particularly high coverage in Kāneʻohe Bay compared to BATS (18.4x vs 0.3x ave. coverage; Supplementary Data 14) that was not found in SAG GCA_902533445.1, and encoded genes for type IV pilus assembly (Supplementary Data 15). The role of type IV pili in SAR11 is unclear^53^, although in other organisms they have been associated with an array of functions including DNA uptake, twitching motility, and aggregation into microcolonies^54^. The presence of type IV pilus assembly genes in the Ia.3.VI genome sourced from the nitrogen-limited Pacific Ocean, but not in genomes from relatively more nitrogen-replete waters of BATS, along with evidence that SAR11 can utilize purine nucleosides and other purine derivatives for nitrogen^11,55^, suggests that the type IV pilus may be advantageous for DNA uptake in nitrogen-poor environments and that HIMB1556 may be locally-adapted.

Contrary to the hypothesis that SAR11 recombines at a rate sufficient to limit ecotypic differentiation within what we have defined as genus-level clusters of SAR11 genomes^47^, our read mapping instead shows that the HIMB1556 and SAG GCA_902533445.1 genomes within cluster Ia.3.VI have sufficiently diverged at the nucleotide level to reveal clear biogeographic divergence, and that they possess sets of genes that reside in hypervariable genomic regions that are clearly associated with the differences in abundance.

We examined gANI estimates, phylogenetic branching, environmental distributions, and ecologically-relevant gene content to support the characterization of ecological diversification at the finest tips of the SAR11 tree, a process that we theorize to represent speciation. This underscores the complexity of SAR11 ecology, highlights the need to include a diversity of representative genomes within even closely related genera for environmental genomics studies, and indicates that continued efforts to sample SAR11 globally are key to understanding the distribution of this ubiquitous clade.

### Proposed *Pelagibacterales* classification and nomenclature

We leveraged the robust genome phylogeny, gANI metrics, and read recruitment to establish a rational classification and nomenclature system for the bacterial order *Pelagibacterales*. To provide a framework and vocabulary to discuss groups of SAR11 in a meaningful context, we first defined four family-level monophyletic groups as the *Pelagibacteraceae* (historical subgroups Ia and Ib), *Cosmipelagibacteraceae* (historical subgroup II), *Fontibacteriaceae* (historical subgroup III), and the *Mesopelagibacteraceae* (historical subgroup Ic) (Fig. 4). We focused our efforts primarily on classification within the *Pelagibacteraceae* where the majority of cultured isolates originate. While we did not exhaustively define genera in the three other *Pelagibacterales* families, we incorporated previously named genera and species, and provided taxonomic names for strains previously cultivated at the Hawaiʻi Institute of Marine Biology (HIMB58 and HIMB114)^37^. Within the *Pelagibacteraceae*, we used phylogenomics and ecological data to characterize 24 genera that represent cohesive genetic and ecological clades, and designate type species for each (Supplementary Data 13). The primary aim of these efforts is to ensure that the taxonomic hierarchy for SAR11 provides a useful and tractable reflection of the ecology and genetic diversity within this globally distributed group, and establishes a rational system that future efforts can build upon.

**Figure 4.**
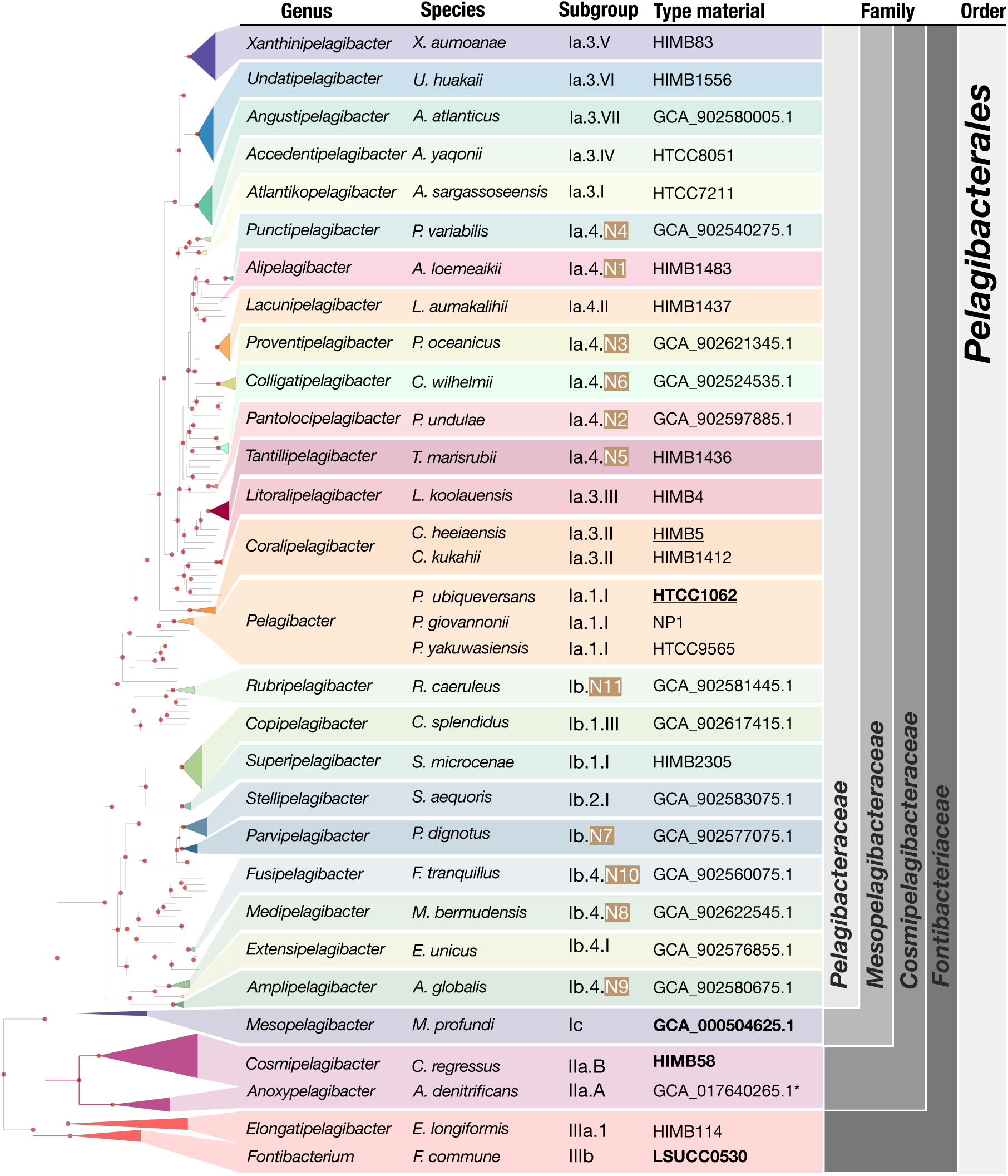
A proposed taxonomic framework for the SAR11 order *Pelagibacterales* that unites proposed genus and species names, type genomes, and historical reference labels. All type genomes for species are presented in the “Type material column”. For instances where there are multiple species characterized within a genus, the designated type species for the genus is underlined. Bold font indicates the type material for each family. The phylogenetic backbone is from the phylogeny generated for Figure 1 and includes branches with genomes that have not yet been classified.

## Discussion

By integrating high-throughput cultivation experiments with publicly available genomes and metagenomes, our study provides key insights into a long-standing question: to what extent, and at what hierarchical levels, can the genomic and ecological diversity of SAR11 be partitioned into cohesive units? Through phylogenomic analyses paired with global metagenomic read recruitment surveys, we reveal ecotypic differentiation at both relatively fine, species-level and coarse, genus-level diversity within SAR11. This robust eco-evolutionary framework, which unifies independent yet complementary approaches to genomic diversity and biogeography, resolves the order *Pelagibacterales* into four families and the family *Pelagibacteraceae* into 24 genera, establishing a much-needed taxonomic framework that delineates SAR11 diversity into tractable units and provides a foundation for future investigations. Such overarching approaches have been shown to be valuable in previous studies^56^, which argued that the absence of a consistent naming structure hinders the identification of coherent ecological traits and emphasized its importance as a foundation for future research.

A tight relationship between the phylogeny and ecology of SAR11 has long been suggested^3,7–10,17,38,57^; however, the ability to associate specific SAR11 clades with distinct ecological patterns and explain forces that maintain SAR11 diversity has remained elusive. Focusing on sequence-discrete groups within deep ocean SAR11 lineages, a recent study concluded that recombination, rather than ecological speciation, was likely the major driver of species-level cohesion^47^. While this observation may explain forces that maintain species-level cohesion for some populations in this group, our study shows that the global sampling of environmental populations through metagenomes consistently supports ecological delineations that are congruent with phylogenomic clustering patterns, pointing towards ecotypic differentiation as the pervasive driver of the evolution within the *Pelagibacterales*. Interestingly, SAR11 genera that showed similar biogeographical distribution patterns in our analysis tended to occupy distant parts of the tree. This observation suggests an inverse correlation between the genetic similarity among SAR11 populations and their co-occurrence, a trend known as phylogenetic overdispersion. Phylogenetic overdispersion has been observed across the tree of life^58^ and is driven by forces of competitive exclusion, an overarching ecological phenomenon that limits the co-occurrence of ecologically similar, closely related organisms. Future analyses that aim to resolve specific genetic determinants of competitive exclusion or co-existence may benefit from geographically constrained time-series data, as these patterns are likely not immediately attainable from global yet spatiotemporally sparse metagenomes. These experimental systems are challenging to maintain, but can be particularly valuable when they document multidimensional aspects of the system that include a range of ʻomics and environmental data^9,11,59–61^.

The practical need of microbiologists to find reasonable cutoffs to demarcate species boundaries from genomic data alone and the nature of SAR11 evolution do not align seamlessly. Through the analysis of genomes, a large number of anecdotal observations support 95% ANI as a reasonable means to resolve archaeal and bacterial species^62,63^. However, SAR11 serves as a reminder that practical solutions do not necessarily apply to all microbial clades^8,10,15,17,47^. One of the implications of the efforts to standardize the tree of life based on principles that work only for the majority of microbial taxa is the conflation of all SAR11 genomes into two genera in the taxonomic framework derived from genomes available on the Genome Taxonomy Database (GTDB) based on RED scores^64^. Indeed, while the ecologically relevant units of SAR11 described in our study are in agreement with functional, evolutionary, and ecological observations, they are in disagreement with the contemporary summaries of this clade based on RED- or ANI-based demarcations. The ways in which evolutionary relationships between distinct clades of life intersect with taxonomic classification systems will unlikely be resolved in a manner that satisfies everyone in microbiology^65,66^. In this juncture, we believe that a stronger motivation to understand the biological drivers that render SAR11 incompatible with our best practical approaches will bring us closer to a unified solution to partition microbial diversity into meaningful units, rather than casting SAR11, one of the most numerous microbial clades on our planet, as a mere outlier.

Insights into the eco-evolutionary processes that shape SAR11 diversification in our study rely heavily on the contribution of 81 new isolate genomes that represent abundant and ecologically-relevant SAR11 populations across the coastal and global ocean. The ecology-informed hierarchical organization of these genomes enabled us to propose SAR11 genera with formal names here. Concurrent efforts from our group investigated the functional determinants of ecological diversification across the *Pelagibacteraceae*^11^ and characterized metabolic capacities and selective pressures that support the evolutionary and ecological partitioning of the genera as proposed here. While deeper understanding of the physiological, metabolic, and genetic factors that shape SAR11 biology will require controlled experimentation of isolated strains in the laboratory, our study organizes the eco-evolutionary characteristics of known SAR11 diversity and provides a roadmap for future efforts aimed to organize and understand the ubiquitous SAR11 populations inhabiting the global ocean.

## Methods

### High-throughput culturing from surface seawater within and adjacent to Kāneʻohe Bay, Oʻahu

Growth medium was prepared following previously described methods^67^. Briefly, 20 L of seawater was collected first on 8 July 2017 and again on 20 September 2017 from a depth of 2 m at station SR4 (N 21° 27.699’, W 157° 47.010’) in acid-washed polycarbonate bottles (Supplementary Fig. 1). The seawater was then filtered, autoclaved, and sparged with CO_2_ and then with air sequentially through three in-line HEPA vent filters (0.3 µm glass fiber, 0.2 µm polytetrafluoroethylene [PTFE], 0.1 µm PTFE; Whatman, GE Healthcare Life Sciences, Chicago, IL, USA), as done previously^67^. After processing, the sterile seawater was stored at 4°C until use.

Two 4 L seawater samples to be used as inoculum were collected on 26 July 2017 in acid-washed polycarbonate bottles from 2 m from stations SB (N 21° 26.181’, W 157° 46.642) and STO1 (N 21° 28.974, W 157° 45.978’) (Supplementary Fig. 1) and immediately returned to the laboratory for further processing. The samples from the Kāneʻohe Bay Time-series (KByT) samples were previously classified as ‘nearshore’, ‘transition’, or ‘offshore’, with SB and STO1 representing nearshore and offshore sites, respectively ^9^. Subsamples of the raw seawater were processed as described previously^67^. Briefly, aliquots were taken for cryopreservation in a final concentration of 10% v/v glycerol and fixed with paraformaldehyde for the enumeration of planktonic microorganisms via flow cytometry. Additionally, 0.96 L from station SB and 1.30 L from station STO1 were filtered through a 25 mm diameter, 0.1 µm pore-sized polyethersulfone membrane (Supor-100; Pall Gelman Inc., Ann Arbor, MI), which was then submerged in 500 µL DNA lysis buffer and stored at -80°C until DNA extraction.

Subsamples of raw seawater from SB and STO1 were enumerated using microscopy, diluted to 2.5 cells mL^-1^, and plated in 2 mL volumes into a total of 1,152 wells (576 wells per site) of custom-fabricated 96-well Teflon microtiter plates. This experiment is referred to here as HTC2017. Plates were then sealed with breathable polypropylene microplate adhesive film and incubated in the dark at 27°C. Plates were monitored for cellular growth at 3.5 and 8 weeks by staining cells with SYBR Green I nucleic acid stain (Invitrogen, Carlsbad, CA, USA) and then counted on a Guava easyCyte 5HT flow cytometer equipped with a 150-mW blue (488-nm) laser (Millipore, Burlington, MA, USA) as done previously^28,67^. Wells with positive growth (greater than 10^4^ cells mL^-1^) after 24 or 57 days of incubation were further sub-cultured by transferring approximately 1 mL into 20 mL of sterile seawater media amended following previous studies^67^ with 400 µM (NH_4_)_2_SO_4_, 400 µM NH_4_Cl, 50 µM NaH_2_PO_4_, 1 µM glycine, 1 µM methionine, 50 µM pyruvate, 800 nM niacin (B3), 425 nM pantothenic acid (B5), 500 nM pyridoxine (B6), 4 nM biotin (B7), 4 nM folic acid (B9), 6 µM myo-inositol, 60 nM 4-aminobenzoic acid, and 6 µM thiamine hydrochloride (B1). These subcultures were then incubated at 27°C in the dark for an additional 33 days and then all samples were processed and cataloged.

Cultures checked at 33 days that yielded positive growth (>10^4^ cells mL^-1^) were cryopreserved in duplicate (2 x 500 µL culture and a final concentration of 10% v/v glycerol). Each well with positive growth was assigned an HIMB culture ID and cells from the approximately 18 mL remaining volume of each culture were collected by filtration through a 13 mm diameter, 0.03 µm pore-sized polyethersulfone membrane (Sterlitech, Kent, WA, USA), which was then submerged in 250 µL DNA lysis buffer and stored at -80°C until DNA extraction. The lysis buffer was prepared by adding the following to Milli-Q water: 8 mL 1M Tris HCl (pH 8.0), 1.6 ml 0.5M EDTA (pH 8.0), and 4.8 g Triton X, for a final volume of 400 mL, which was then filter sterilized, with lysozyme added to aliquots immediately before use (at a final concentration of 20 mg mL^-1^) as done in prior studies^41^.

An additional experiment was performed using cryopreserved samples of seawater collected on July 26, 2017, and described previously^67^. Briefly, the cryopreserved sample was enumerated and then diluted to two cell concentrations (2.5 and 52.5 cells mL^-1^), and used to plate 480 and 470 2-mL dilution cultures, respectively. This experiment is referred to as HTC2018. Growth was monitored at 2, 3, and 5 weeks after inoculation with positive growth (>10^4^ cells mL^-1^) from the 2.5 cells mL^-1^ cultures subcultured into 20 mL of sterile seawater growth medium and monitored for growth for up to 10 weeks at 27°C in the dark. Subcultures were then cryopreserved and cells collected for DNA sequencing as described above. One well from the 52.5 cells mL^-1^ inoculation was directly collected for DNA sequencing without subculturing^67^.

### DNA extraction and 16S rRNA gene amplicon sequencing

Genomic DNA (gDNA) from all filtered cultures as well as environmental DNA (eDNA) from STO1 and SB was extracted using the Qiagen DNeasy Blood and Tissue Kit with modified manufacturer’s instructions for bacterial cells (Qiagen, Germantown, Maryland, USA). The modifications included the addition of an initial freeze-thaw step (3 cycles of 10 minutes at 65°C followed by 10 minutes at -80°C), the addition of 35 µL Proteinase K and 278 µL buffer AL at the appropriate pretreatment step, and finally when eluted the same 200 µL volume was passed through the membrane three times.

For the initial identification of all cultures, gDNA was used as template for the polymerase chain reaction (PCR) amplification (Bio Rad C1000 Touch, Bio Rad, Hercules, CA, USA) using barcoded 515F-Y (5′-GTGYCAGCMGCCGCGGTAA-3′) and 926R (5′-CCGYCAATTYMTTTRAGTTT-3′) primers targeting the V4-V5 region of the SSU rRNA gene^68^ in a reaction volume of 25 µL composed of: 2 µL gDNA, 0.5 µL each forward and reverse primer, 10 µL 5PRIME HotMasterMix (Quantabio, Beverly, MA, USA), and 12 µL of molecular grade H_2_0^67^. The reaction was as follows: an initial denaturation step of 3 min at 94°C, 40 cycles of 45 sec at 94°C followed by 1 min at 50°C and 1.5 min at 72°C, with a final extension of 10 min at 72°C. The PCR products were quantified, pooled at a concentration of 240 ng per sample, and cleaned (QIAquick PCR purification kit; Qiagen) prior to sequencing following previous methods^67^ and sequenced on a MiSeq platform by the Oregon State University Center for Genome Research and Biocomputing.

### 16S rRNA gene sequence analysis

Amplicon sequence data were processed following previous methods^67^. Briefly, the data was imported into QIIME2 v2019.4.0, and demultiplexed before being assessed for sequence quality and merged. DADA2 v1.14^69^ was then used for quality control. Taxonomy was assigned to all reads using a Naïve Bayes classifier trained on the Silva rRNA v132 database^70^. Cultures were first classified as defined previously^67^, with “monocultures’’ consisting of more than 90% of reads from a single amplicon sequence variant (ASV), “mixed cultures’’ with an ASV that was between 50% and 90% of the reads, and finally cultures with no dominant members. Any samples with less than 1,000 reads were not included in further analyses. We aimed to sequence all strains that included monocultures and mixed cultures of SAR11.

### Genome sequencing

For all extractions with gDNA concentrations >0.06 ng µL^-1^, 10 µL were aliquoted for sequencing. For samples with concentrations <0.06 ng µL^-1^, the remaining extraction volume (∼180 µL) was concentrated using a SpeedVac (ThermoFisher) to approximately 30 µL and re-quantified (Qubit 2.0, Invitrogen). Concentrated samples with a minimum of 0.06 ng µL^-1^ were then aliquoted (10 µL) for sequencing. Samples for sequencing were prepared using a Nextera library kit and sequenced on the NextSeq500 platform via a 150 bp paired-end run.

Genomes for previously cultured strains HIMB109 and HIMB123^37^ were sequenced by the US Department of Energy’s Joint Genome Institute. Multiple methods were used to sequence the two strains, including directly using 200 µL of cell culture for library prep as well as using multiple volumes (5, 10, or 20 µL) of culture for multiple displacement analysis (MDA) prior to library preparation (Supplementary Methods). The two genomes were evaluated based on completeness, length, number of reads, and total contigs post-assembly using SPAdes 3.12.0^71^. An additional assembly using all reads generated from the various sequencing attempts from each strain was also constructed using the same assembly method. Based on the metrics described above, the highest quality genomes were manually curated and used for additional analyses.

### Genome assembly and assessment

Short reads were trimmed with Trim Galore! (https://github.com/FelixKrueger/TrimGalore) and assembled using Unicycler v0.4.8^72^, which acts as a SPAdes 3.13.0^71^ optimizer with Illumina short read data. Once assembled, reference indexes were built, and read mapping was performed using Bowtie2 with default parameters^73^. SAMtools^74^ was used to convert the SAM file to a sorted and indexed BAM file. These initial assemblies and BAM files were used to visualize genomes in anvi’o v8.0 to check for possible contamination^75,76^. For genomes with contamination (determined visually as instances where contigs had anomalous GC content or tetranucleotide frequency) suspicious contigs were removed. Redundancy was also used as a way to flag any genomes that needed further curation. After inspection, curated contigs were exported using the program ‘anvi-summariz’ and reads were re-mapped to the cleaned version of assemblies. The cleaned genomes were processed again for visualization in anvi’o to ensure no erroneous contigs were included. Mapping quality was inspected visually using the Integrative Genomics Viewer (IGV) v2.8.4^77^ and Tablet v1.21.02.08^78^ and manual curation was undertaken using mapped read data. Manual inspection was used to determine if a circular genome could be considered closed and complete. All contigs shorter than 1000 bp were removed from the genomes that were not closed after final curation, and CheckM 1.1.2 was used to assess final genome completeness and redundancy^79^.

### Phylogenomic analyses

To generate an extended phylogeny of the SAR11 clade, a suite of high-quality genomes were curated. Even with an abundance of metagenomes, the high diversity among SAR11 populations makes constructing reliable MAGs currently unfeasible^20^, so to ensure the phylogeny was as robust as possible, only isolate genomes and SAGs were included. The final set of 484 SAR11 genomes for phylogenetic reconstruction included 81 genomes sequenced in this study, 28 previously published reference genomes, and 375 previously published single amplified genomes (SAGs), in addition to 10 isolate genomes from the family *Rhodobacteraceae* used as an outgroup (Supplementary Data 3). The majority of SAGs included were equal to or greater than 85% complete according to CheckM 1.1.2^79^. However, genomes of lower quality from subclades of SAR11 with no high-quality representatives were included to produce a phylogeny that best represents established clades, for example SAR11 Ic genomes that ranged from 56.0 to 93.7% completion were also included^39^ (Supplementary Data 3). For comparison, we also constructed a phylogeny including 27 available genomes from what were previously SAR11 subgroups V and IV. Subgroups V and IV are no longer considered to be within SAR11 and the membership of subgroup IV has not been rigorously investigated, thus its relationship to the *Pelagibacterales* is questionable as we further describe in the Supplementary Notes^10,22,43,45^.

We compared two sets of phylogenetic markers to determine the most appropriate genes to use for phylogenetic reconstructions of the SAR11 clade. This included the proteins defined by the bac120 set utilized by GTDB-Tk to determine the bacteria guide tree, and a curated set of markers derived from the 200-genes previously demonstrated to be best fit for the *Alphaproteobacteria*^23^ (Supplementary Data 4). An alignment file of the proteins based on the bac120 set was generated using GTDB-Tk^80^. To curate the second set of phylogenetic markers, we generated a custom HMM profile for the 200 *Alphaproteobacteria* genes with a noise cutoff term of 1×10^−20^ , ran the HMM profile on all genomes using the anvi’o v8.0 program ‘anvi-run-hmms`’`, and generated a presence-absence matrix of genes in this model across genomes using the program ‘anvi-script-gen-hmm-hits-matrix-across-genomes`’`. After evaluating the model hits across the genomes matrix, we removed the genes that occurred in less than 90% of the genomes or those that were redundant in more than 2% of the genomes from the *Alphaproteobacteria* 200-gene collection, which resulted in a new collection with 165 genes, which is referred to as the ‘SAR11_165’ throughout our study (Supplementary Data 4). To generate a concatenated alignment of the proteins of interest for downstream phylogenomic analyses, a custom HMM source was generated that encompassed the SAR11_165 marker protein sequences. The program ‘anvi-get-sequences-for-hmm-hits`’` with the custom HMM source was then implemented to extract and align proteins of interest, which uses Muscle (v3.8.1551) to align sequences^81^. For the alignments from each set, the program trimAL 1.3^82^ was then used to remove all positions that were missing in more than 50% of the genomes. Phylogenies were generated with IQ-Tree v2.1.2^83^ with the best fit model (LG+F+R10) chosen using ModelFinder^84^ and 1,000 ultrafast bootstraps. Phylogenies were rerooted appropriately in FigTree v1.4.4 using the midpoint rooting method, and exported in NEXUS format with the options selected to “Save as currently displayed” and “Include Annotations (NEXUS & JSON only)”. Once exported, phylogenies were then compared using the package phytools v1.2-0^85^ in R v4.2.1^86^.

Once the phylogeny was established, a subset of SAR11 genomes was used for read recruitment. For this, we first used FastANI v1.34^62^ to dereplicate all genomes using 95% gANI as a cutoff, then for genomes within the historical Ia and Ib clades, we excluded SAGs that did not share at least 90% gANI with a neighboring genome. A 95% gANI dereplication cutoff was selected to avoid read splitting during competitive read recruitment and for any clusters in which isolate genomes were available, they were chosen as preferred representatives.

### Classification and nomenclature

The phylogeny was used to define cohesive genetic clusters at the distal end of the SAR11 tree. Single genomes that did not share at least 90% gANI with a neighboring genome in the historical Ia or Ib subclades were not classified into genera.

To determine how taxonomic levels across the SAR11 lineage would compare using relative evolutionary distance (RED) values, we implemented this approach following previous studies^59^ (Supplementary Notes). Briefly, a domain-level phylogeny was first constructed using the GTDB-Tk de_novo_workflow^80^ with SAR11 isolate genomes and SAGs as well as “p Chloroflexota” as the outgroup. Marker genes were identified from the input genomes using GTDB-Tk ‘identify`’`, and then aligned with GTDB-Tk ‘align`’` (using the “–skip_gtdb_refs” flag). Finally, a tree was constructed using FastTree v2.1.10 (model WAG+GAMMA)^87^, rooted with the Chloroflexota outgroup. This phylogeny was used as the input for the ’scale_tree’ program in PhyloRank v0.1.11 (https://github.com/dparks1134/PhyloRank) to convert branch lengths into RED scores. RED values of 0.77 and 0.92 were used to assess how they would align with family and genus-level lineages, respectively. These values were based on the distribution of internal nodes within the SAR11 clade and values used previously for other family and genus-level lineages^88^.

To provide species names that utilize descriptive ʻŌlelo Hawaiʻi (the Hawaiian language) words for species with type material that was cultivated on the island of Moku o Loʻe located in Kāneʻohe Bay in the watershed of Heʻeia, we collaborated with local community leaders. Guided by collaborative research protocols practiced in Heʻeia^89,90^, we received input from Kumu Hula (Master Instructor), Frank Kawaikapuokalani Hewett and coordinated with the Indigenous Stewardship Specialist and Stewardship Coordinator at the Heʻeia National Estuarine Research Reserve, Aimee Saito.

### Read recruitment

To assess the distribution of the newly described strains described in this study and put them into context with previously sequenced genomes, we used a read recruitment approach with globally distributed metagenomes. The SAR11 genomes included in this study were grouped into clusters that shared 95% genome-wide average nucleotide identity (gANI) or greater (Supplementary Data 5) and representatives from these 95% gANI groups were then used for read recruitment to ensure reads were not split between closely related genomes. Results from read recruitment were extrapolated for the other genomes included in each 95% gANI group.

Metagenomes used for recruitment included those sequenced in Kāneʻohe Bay^60^, the environment from which the genomes were isolated. Only samples from sites previously categorized as “nearshore” and “offshore”^60^ were used here. Additionally, globally distributed previously published metagenomes were also used including those from TARA Oceans expeditions^91^, Station ALOHA^92^, GEOTRACERS cruises^93^, the eastern coast of Japan^94,95^, Monterey Bay^96^, and the Ocean Sampling Day (OSD) program^97^ (Supplementary Data 7 for a list of appropriate references and details regarding metagenomes included).

Once metagenomes were chosen, raw reads were downloaded using ‘prefetch’ and ’fasterq-dump’ in the SRA toolkit v2.10.1. We automated the quality filtering of metagenomes, metagenomic read recruitment, and profiling of recruited reads using the program anvi-run-workflow^98^ with the ‘--workflow metagenomics’ flag, which implements snakemake^99^ recipes for standard analyses in anvi’o. Briefly, this workflow identified and discarded the noisy sequencing reads in metagenomes using the program ‘iu-filter-quality-minoch’^100^, used SAR11 genomes to competitively recruit short reads from metagenomes using Bowtie2^73^ SAMtools^74^ using the program ‘anvi-profile`’`, and finally merged individual profiles into an anvi’o merged profile database using the program ‘anvi-merg ’. The resulting anvi’o merged profile database included essential data, including genome coverages and detection statistics across metagenomes, for our downstream analyses. For coverage, we primarily used the ‘mean coverage Q2Q3’ statistic, which represents the interquartile average of coverage values where, for any given genome, the lowest 25% and the highest 25% of individual coverage values are trimmed prior to calculating the average coverage from the remaining data points, and thus minimizing the impact of biases due to highly conserved or highly variable regions in the final coverage estimates. The detection metric equated to the total proportion of nucleotides in a given contig that are covered at least 1X, we used the previously established threshold of 0.25 to state that a given genome was present in a sample and remove any potential false positives^101^. Visualization of read recruitment data mapped according to the phylogeny constructed was completed using the program ‘anvi-interactive`’` with the ‘--manual`’` flag.

### Metagenome profile clustering

We performed a cluster analysis of metagenomes based on genome detection values from the read recruitment step using the k-means algorithm, where we determined the ‘k`’` by identifying the elbow of the curve of within-cluster sum of square values for increasing values of ‘k’ using the R code shared by Delmont et al.^8^ at https://merenlab.org/data/sar11-saavs/. The results of the clustering analysis were visualized using anvi’o and the vegan package v2.6-4 in R v4.2.1^102^ was used to perform a permutational multivariate analysis of variance (PERMANOVA) to test the significance of the resulting groupings using the Bray-Curtis method of calculating the dissimilarity matrix and the adonis2 function. This method was also used to test the significance of genomes grouped by subclade. Additionally, the “betadisper” function in vegan was used to conduct a PERMDISP, which was visualized via a principal coordinates analysis (PCoA). To investigate how similar the detection patterns of genomes within and between genome clusters were, we performed a non-metric multidimensional scaling (NMDS) analysis using the vegan package v2.6-4 in R. Any metagenomes with zero detection across all genomes were removed. The NMDS results were visualized using ggplot2 v2_3.5.1 and plotly v4.10.4 and an interactive plot was generated with ggplotify v0.1.2.

### Investigation of C-P lyase pathway

All SAR11 genomes included in the extended phylogeny (n=481) were searched using ‘anvi-search-functions’ for the key enzyme in the C-P lyase pathway (*phnJ*) to determine the capacity among high-quality SAR11 genomes to utilize the pathway. The genes upstream and downstream of this essential gene were extracted from all 57 genomes carrying this gene and a pangenome was used to compare the presence and absence of other key genes in the pathway as well as the synteny of this region of the genome. A phylogeny was constructed from an alignment of PhnJ protein sequences recovered from the genomes (Supplementary Fig. 7).

## Data availability

We have deposited the assembled sequence data for genomes, including raw sequencing reads, under the NCBI BioProject PRJNA1170004. The Supplementary Data include all remaining data, and the URL https://seqco.de/r:r4auejub serves as the SeqCode registry for all taxon names defined in this study.

## Code availability

All custom R, BASH, and Python scripts used for data analyses in this study are publicly available on GitHub at https://github.com/kcfreel/SAR11-genomes-from-the-tropical-Pacific and archived with Zenodo (https://doi.org/10.5281/zenodo.17614008). Additionally, a fully reproducible bioinformatics workflow for the analysis of SAR11 genomes is available at https://merenlab.org/data/sar11-phylogenomics/, enabling the reproduction of our phylogenomic tree and its extension with new genomes.

## Acknowledgments

We thank Kumu Hula, Kawaikapuokalani Hewett and Aimee Sato for their generous guidance in using approprate ʻŌlelo Hawaiʻi (Hawaiian Language) words to determine species names of isolates cultivated on Moku o Loʻe. We also thank K. Luttrell for help with Latin grammar, R. Malmstrom and N. Nath for sequencing the genomes of isolates HIMB109 and HIMB123, R. Ouye for assistance with HTC experiments, and O. Ramfelt for bioinformatic support. We also thank F. Trigodet for their help with the high-performance computing at the University of Oldenburg. Finally, we sincerely thank Luis Miguel Rodriquez-R, Marike Palmer, and the entire SeqCode team for their expert grammatical and taxonomic guidance. This research was supported by funding from the National Science Foundation grants OCE-1538628 (MSR) and OCE-2149128 (MSR) as well as the Simons Postdoctoral Fellowship in Marine Microbial Ecology (LS-FMME-00989028) (SJT).

## Author contributions

KCF, SJT, AME, MSR conceived the study, developed methodology, and led the investigation and visualization for the study. AME and MSR supervised the study. KCF wrote the original draft. KCF, SJT, EBF, US, SJG, AME, MSR reviewed and edited the manuscript.

