## Supplementary information for "New SAR11 isolate genomes and global marine metagenomes resolve ecologically relevant units within the *Pelagibacterales*"

Kelle C. Freel *et al*.

**This PDF file includes:**

Supplementary Methods
Supplementary Notes
Supplementary Fig. 1 to Supplementary Fig. 10
Supplementary References

#### Supplementary Methods

##### Genome sequencing of HIMB109 and HIMB123 at the Joint Genome Institute (JGI)

A Nextera XT library prep kit was used to process samples before sequencing on a NextSeq platform (Illumina). Raw data was processed with BBDuk (version 37.36, BBMap – Bushnell B. – sourceforge.net/projects/bbmap/) to remove contamination, trim reads with adapter sequences, and trim the ends of reads with quality scores below 6. This step also removed reads with 2 or more 'N's', less than or equal to 49 base pairs in length, and with an average quality score across the read less than 10. Reads mapped with BBMap (BBMap – Bushnell B. – sourceforge.net/projects/bbmap/) to masked human, cat, dog, and mouse references at 93% identity were separated into a separate file. Reads that aligned to common microbial contaminants [JGI SOP 1077] were also split into a separate file.

##### 16S rRNA gene amplicon sequence analysis

The 16S rRNA gene sequence analysis was previously described[1]. Briefly, three Illumina MiSeq runs were imported into QIIME2 v2019.4.0 and demultiplexed, and paired ends were analyzed for sequence quality and merged[2]. The DADA2 software package[3] was used to denoise sequences, which included removing chimeras. Due to the low quality at the end of the sequences, 10 bases were truncated from the 3′ end of the reverse reads. Sequence reads from the three runs were then merged post-denoising. Amplicon sequence variant (ASV) identities were defined by DADA2 for all reads that varied by at least 1 bp. Taxonomy was assigned to each ASV using a Naïve Bayes classifier trained on the Silva rRNA v132 database[4], clustered at 99% similarity, and subsequently modified manually based on phylogenetic analyses and the results of previous work.

The ASV identities were further evaluated in QIIME2, where any ASVs represented by less than 20 reads were removed from the dataset. The proportion of each ASV in an individual culture was determined by dividing the read count for that ASV by the total number of reads from the culture, post-curation. Cultures were identified as (1) monocultures (dominant member $\geq$90% of reads), (2) mixed cultures (dominant member $\leq$90% of reads and an additional member $\geq$5% of reads), or (3) cultures with no dominant member (no ASV $\geq$50% of reads). Cultures in the third category were not included in additional analyses.

##### Reference genomes

A literature search was used to identify publicly available isolate genomes and high-quality single amplified genomes (SAGs) (Supplementary Data 3). Although the majority of the included SAGs were derived from a single study[5], the samples used for sequencing were globally distributed. The site from which each genome originated is included in Supplementary Data 3 for all genomes except for HTCC9022, for which no specific latitude or longitude was available. However, it was confirmed that it originated from ocean surface waters, offshore, over the Juan De Fuca Ridge. To include a diversity of SAGs and ensure genomes were of reliable quality, we used an 85% completion cutoff for inclusion in this study, except for cases where an established SAR11 lineage had limited or no genomes available that met these criteria. Additional SAGs were included from the Mediterranean Sea[6], those representing the deep SAR11 Ic clade[7], genomes from oxygen minimum zones[8], and the Red Sea[9]. To ensure no high-quality SAGs were excluded from the analyses, we cross-referenced the 'hq_mimag_genomes' file for the release of the Genome Taxonomy Database (GTDB) from April 23, 2024. This file is the complete list of isolate, MAG, and SAG genomes that meet the MIMAG high-quality genome criteria[10] which includes completeness values greater than 90% and contamination less than 5% in addition to the presence of 5S, 16S, and 23S rRNA genes with the minimum lengths of 80 bp, 1200 bp, and 1900 bp, respectively for bacterial genomes. Additionally, these genomes all encode a minimum of 18 tRNAs. For all genomes classified in GTDB as "o__Pelagibacterales", there were a total of 130 high-quality genomes, 11 of which were MAGs (including 6 from freshwater lake metagenomes), and 91 mislabeled as MAGs but instead are SAGs derived from the study of [5].

The type material for 'Mesopelagibacter carboxydoxydans' and *Anoxypelagibacter denitrificans* are MAGs ETNP_2013_S10_SV74_125m_MAG_02 (GCA_017640275.1) and ENTP2013_S02_SV82_300m_MAG_01 (GCA_017640265.1), respectively[11]. To maintain dataset consistency and avoid the population-level heterogeneity and contamination that accompanies SAR11 MAGs[12], we used high-quality single amplified genomes as representatives of these sublineages in our analyses. To verify the relationship between the designated type material MAGs and substitute SAGs, we constructed phylogenies with the SAR11_165 gene set from the Ic and II subclades, respectively. In both cases, 10 isolate genomes from the *Rhodobacteraceae* were included as an outgroup. To generate a concatenated alignment, the program 'anvi-get-sequences-for-hmm-hits' with the custom HMM source was implemented to extract and align genes of interest. For the alignments from each gene set, the program trimAL 1.3[13] was then used to remove all positions missing in more than 50% of the genomes. Phylogenies were generated with IQ-Tree v2.1.2[14] with the LG+F+R10 model and 1,000 ultrafast bootstraps.

**Figure maps**

The Hawai'i, O'ahu, and the larger global maps were made in R with the following packages: dplyr v1.1.4, stringr v1.5.2, ggplot2 v4.0.0, maps v.3.4.3, sf v1.0-22, marmap v1.0.12, mapdata v2.3.1, and ggspatial v1.1.10. Images were then saved and imported to Affinity Publisher v1.10.8 (Figure 3d) or Affinity Design v1.10.5 (Supplementary Figure 1) for finalization. The coastal map of Kāne'ohe Bay has bathymetry lines traced from Tucker et al., (2025)[28], and the base map was generated in ArcGIS for Monagahan et al., (2020)[29].

#### Supplementary Notes

**Cultured isolates represent dominant and persistent SAR11 populations in Kāne'ohe Bay**

We investigated the ecological relevance of the recently isolated strains by recruiting metagenomic reads from a three-year time series of coastal and offshore tropical marine plankton collected as a part of the Kāne'ohe Bay Time-series (Supplementary Fig.1; Supplementary Fig. 8), as well as public sequence data from microbial communities at open-ocean Station ALOHA in the adjacent North Pacific Subtropical Gyre (NPSG). Within the tropical Pacific, fifteen genomes from cultivated HIMB strains are among the limited number of *Pelagibacterales* genomes persistently detected nearshore (Supplementary Fig. 8). Six of these genomes were sequenced from cultures first grown in the laboratory in 2005[15], and their detection in metagenomes from samples collected more than 10 years later (2017 through 2019) underscores the temporal persistence of cultivated populations of SAR11 in Kāne'ohe Bay. The dominant and persistent environmental SAR11 populations are well represented by isolate genomes, and the nearshore environment within Kāne'ohe Bay retains limited SAR11 diversity compared to the adjacent open ocean.

**Relative evolutionary distances do not define meaningful units of SAR11 diversity**

We examined how the taxonomic classification system proposed here compares to the taxonomic groupings of GTDB and those resulting from the common methodology of Relative Evolutionary Distance (RED) (Supplementary Fig. 9, Supplementary Data 16). The three classification systems (GTDB, RED values, and the one proposed here) reported non-congruent groupings of SAR11 diversity. The GTDB classification system results in polyphyletic clusters at the levels of family, genus, and species when overlaid on our phylogenomic tree. To assess the RED approach, we compared the results of categorizing the genomes using RED value thresholds that in other microbial lineages roughly correspond to family (0.77), genus (0.88), and species levels (0.92) of classification. Using RED values, we found instances of both under- and over-splitting in the family- and genus-level designations compared to our proposed taxonomic framework. For example, the RED value of 0.77 leads to a clade encompassing all SAR11 subclade Ic, Ib, and Ia genomes into a single family, whereas we delineate two families through our ecological and evolutionarily-guided analyses. Furthermore, implementing a RED value of 0.92 placed all Ia.3 and Ia.4 sublineages together even though they exhibit distinct spatiotemporal patterns of distribution, contain well-supported branching patterns concomitant with distinct genome properties such as genome size and GC content, experience variable selective pressure[16], and harbor variable gene content associated to distinct metabolic characteristics[16]. While RED values can be crucial tools to reflect meaningful classification schemes[17], including for the enigmatic SAR86 clade[18], this metric fails to distinguish meaningful units of diversity within SAR11.

**Relationship between 'Mesopelagibacter carboxydoxydans' and *Anoxypelagibacter denitrificans* type material and SAGs of the curated dataset**

To compare the SAGs included in our analyses with the MAGs designated as type material for 'Mesopelagibacter carboxydoxydans' and *Anoxypelagibacter denitrificans,* we generated phylogenies that included the type genomes, the related SAGs included in our dataset, and the *Rhodobacteraceae* outgroup. The type material for 'Mesopelagibacter carboxydoxydans' (GCA_017640275.1) and *Anoxypelagibacter denitrificans* (GCA_017640265.1) are most closely related to GCA_902582665.1 (88.6% gANI) and GCA_902578935.1 (79.7% gANI), respectively. In both cases, the type material genomes originate from within the monophyletic clades that include the SAG genomes used in our study (Supplementary Fig. 10).

**Other taxonomic considerations of note**

In order to validly publish our taxonomic framework for SAR11 through the SeqCode [19], some amendments to existing taxa were needed. First, the initially proposed species name for the type material of the order, '*Pelagibacter ubique*'[20], has long been known to be invalid. While other options have been proposed [e.g. 'communis'[21]], we sought a species name that retained a linguistic connection to the original and so amended the nomenclatural type from '*Pelagibacter ubique*' to *Pelagibacter ubiqueversans.*

The type material for 'Mesopelagibacter carboxydoxydans', MAG ETNP_2013_S10_SV74_125m_MAG_02 (GCA_017640275.1), is of insufficient quality to be used as type material within the SeqCode. We sought to retain the genus name as proposed by the authors of the effective publication[11] and so designated a new species, *Mesopelagibacter profundi*, that includes a high quality SAG (GCA_000504625.1) as type material. This allowed us to then propagate this genus name to the family-level lineage *Mesopelagibacteraceae* to encompass all of SAR11 subclade Ic.

#### Protologues

##### Description of *Mesopelagibacteraceae* fam. nov.

*Mesopelagibacteraceae* ([Me.so.pe.la.gi.bac.te.ra'ce.ae] N.L. masc. n. *Mesopelagibacter*, the type genus of the family; L. fem. pl. suff. -*aceae*, ending to denote a family; N.L. fem. pl. n. *Mesopelagibacteraceae*, the *Mesopelagibacter* family).

The family *Mesopelagibacteraceae* encompasses what has previously been characterized as the monophyletic sublineage Ic of SAR11 marine bacteria[7,11,22]. The members of this family are free-living cells that predominantly reside in the mesopelagic region of the open ocean. The type genus is *Mesopelagibacter*.

Registry URL: https://seqco.de/i:51441

##### Description of *Cosmipelagibacteraceae* fam. nov.

*Cosmipelagibacteraceae* ([Cos.mi.pe.la.gi.bac.te.ra'ce.ae] N.L. masc. n. *Cosmipelagibacter*, the type genus of the family; L. fem. pl. suff. -*aceae*, ending to denote a family; N.L. fem. pl. n. *Cosmipelagibacteraceae*, the *Cosmipelagibacter* family).

The family *Cosmipelagibacteraceae* encompasses what has previously been described as SAR11 subclade II[23]. The type genus *Cosmipelagibacter* contains the only strain currently isolated from this family, HIMB58[15,24]. One other previously described genus (Anoxypelagibacter) is also included within the family *Cosmipelagibacteraceae*[11].

Registry URL: https://seqco.de/i:51442

##### Description of *Fontibacteriaceae* fam. nov.

*Fontibacteriaceae* ([Fon.ti.bac.te.ri.a'ce.ae] N.L. neut. n. *Fontibacterium*, the type genus of the family; L. fem. pl. suff. -*aceae*, ending to denote a family; N.L. fem. pl. n. *Fontibacteriaceae*, the *Fontibacterium* family).

The family *Fontibacteriaceae* encompasses previously described SAR11 subclades IIIa and IIIb, which have also been referred to as LD12 or subgroup IV[25]. To note, subgroup IV of Morris et al., (2005)[25] does not correspond to subclade IV. The family currently circumscribes two genera that contain strains isolated from either freshwater environments (type genus *Fontibacterium*[26], or surface seawater (*Elongatipelagibacter*, newly described here).

Registry URL: https://seqco.de/i:51443

##### Emendation to the description of the family *Pelagibacteraceae*

We amend the description of the family *Pelagibacteraceae* to encompass historical subclades Ia and Ib of the SAR11 lineage. Previously, the family *Pelagibacteraceae* was described to encompass the entire SAR11 lineage[27]. The type genus remains *Pelagibacter*.

##### Emendation to the description of the genus *Pelagibacter*

We amend the description of the genus *Pelagibacter* to include the type species, *P. ubiqueversans* str. HTCC1062$^{T}$ [20], and its closely related monophyletic relatives within the sublineage previously recognized as SAR11 Ia.1.I. Previously, the genus *Pelagibacter* encompassed the entire SAR11 Ia sublineage[24].

**Emendation to the description of the genus *Mesopelagibacter***

We amend the description of the genus *Mesopelagibacter* to include the type species, *Mesopelagibacter profundi*. The genome of the previously proposed type species, 'Mesopelagibacter carboxydoxydans', could not be validated because the type genome was of insufficient quality (MAG ETNP_2013_S10_SV74_125m_MAG_02; GCA_017640275.1)[11]. The type genome of *Mesopelagibacter profundi* is the high quality SAG GCA_000504625.1.

Registry URL: https://seqco.de/i:51491

**Description of *Xanthinipelagibacter* gen. nov.**

*Xanthinipelagibacter* ([Xan.thi.ni.pe.la.gi.bac'ter] N.L. fem. n. *xanthina*, xanthine, a purine base; L. neut. n. *pelagus*, the open sea; N.L. masc. n. *bacter*, a rod; N.L. masc. n. *Xanthinipelagibacter*, a rod shaped bacterium from the open sea that can metabolize xanthine).

Genomes of the genus *Xanthinipelagibacter* occupy a monophyletic node within the family *Pelagibacteraceae* that is currently known as SAR11 subclade Ia.3.V, and share a minimum gANI of ~85% to the type genome. Cells of this genus contain the genetic machinery for metabolizing xanthine[16], and have been isolated from the tropical Pacific Ocean and Tannan Bay, China. The type species is *Xanthinipelagibacter aumoanae*.

Registry URL: https://seqco.de/i:51392

**Description of *Undatipelagibacter* gen. nov.**

*Undatipelagibacter* ([Un.da.ti.pe.la.gi.bac'ter] L. masc. perf. part. *undatus*, surged, risen in waves, flooded, welled up; L. neut. n. *pelagus*, the open sea; N.L. masc. n. *bacter*, a rod; N.L. masc. n. *Undatipelagibacter*, a surging oceanic bacterium, referring both to the oceanic habitat and to the comparatively large genome size found in this lineage).

Genomes of the genus *Undatipelagibacter* occupy a monophyletic node within the family *Pelagibacteraceae* that is currently known as SAR11 subclade Ia.3.VI, and share a minimum gANI of ~88% to the type genome. This group harbors the largest mean genome size compared to other genera of the*Pelagibacteraceae* genera (~1.43 Mbp). Cells of this genus have been isolated from the tropical Pacific Ocean. The type species is *Undatipelagibacter huakaii*.

Registry URL: https://seqco.de/i:51394

**Description of *Angustipelagibacter* gen. nov.**

*Angustipelagibacter* ([An.gus.ti.pe.la.gi.bac'ter] L. neut. n. *angustum*, a narrow place; L. neut. n. *pelagus*, the open sea; N.L. masc. n. *bacter*, a rod; N.L. masc. n. *Angustipelagibacter*, a rod from a narrow place in the sea, referring to the limited geographic locations from which most genomes originate).

Genomes of the genus *Angustipelagibacter* occupy a monophyletic node within the family *Pelagibacteraceae* that is currently known as SAR11 subclade Ia.3.VII, and share a minimum gANI of ~86% to the type genome. There are no known cultivated isolates reported for this genus; the majority of publicly available single amplified genomes (SAGs) have been recovered from the Mediterranean Sea[6] and Atlantic Ocean[5]. The type species is *Angustipelagibacter atlanticus.*

Registry URL: https://seqco.de/i:51396

**Description of *Atlantikopelagibacter* gen. nov.**

*Atlantikopelagibacter* ([At.lan.ti.ko.pe.la.gi.bac'ter] Gr. masc. adj. *atlantikos*, Atlantic; L. neut. n. *pelagus*, the open sea; N.L. masc. n. *bacter*, a rod; N.L. masc. n. *Atlantikopelagibacter*, bacteria of the open sea from the Atlantic ocean).

Genomes of the genus *Atlantikopelagibacter* occupy a monophyletic node within the family *Pelagibacteraceae* that is currently known as SAR11 subclade Ia.3.I, and share a minimum gANI of ~92% to the type genome. Strains of this genus have been isolated from the site of the Bermuda Atlantic Time-series Study in the Sargasso Sea of the Atlantic Ocean. The type species is *Atlantikopelagibacter sargassoseensis.*

Registry URL: https://seqco.de/i:49683

**Description of *Accedentipelagibacter* gen. nov.**

*Accedentipelagibacter* ([Ac.ce.den.ti.pe.la.gi.bac'ter] L. masc. pres. part. *accedentus*, approaching; L. neut. n. *pelagus*, the open sea; N.L. masc. n. *bacter*, a rod; N.L. masc. n. *Accedentipelagibacter*, the approaching oceanic bacterium, referring to the identification of this bacterium close to the coast in Newport, Oregon).

Genomes of the genus *Accedentipelagibacter* occupy a monophyletic node within the family *Pelagibacteraceae* that is currently known as SAR11 subclade Ia.3.IV, and share a minimum gANI of ~86% to the type genome. Cells of this genus have been isolated from the temperate northeast Pacific Ocean. The type species is *Accedentipelagibacter yaqonii.*

Registry URL: https://seqco.de/i:51398

**Description of *Colligatipelagibacter* gen. nov.**

*Colligatipelagibacter* ([Co.lli.ga.ti.pe.la.gi.bac'ter] L. masc. perf. part. *colligatus*, bound together; L. neut. n. *pelagus*, the open sea; N.L. masc. n. *bacter*, a rod; N.L. masc. n. *Colligatipelagibacter*, a rod bound together across the open sea, referring to the cohesive distribution of this genus).

Genomes of the genus *Colligatipelagibacter* occupy a previously uncharacterized monophyletic node within the Ia.4 cluster of the family *Pelagibacteraceae*, and share a minimum gANI of ~91% to the type genome. There are no known cultivated isolates yet reported for this genus. The type species is *Colligatipelagibacter wilhelmii.*

Registry URL: https://seqco.de/i:51400

**Description of *Tantillipelagibacter* gen. nov.**

*Tantillipelagibacter* ([Tan.ti.lli.pe.la.gi.bac'ter] L. masc. adj. *tantillus*, very small and little; L. neut. n. *pelagus*, the open sea; N.L. masc. n. *bacter*, a rod; N.L. masc. n. *Tantillipelagibacter*, a very small oceanic bacterium, referring to the small genome size of members of this group compared to close relatives).

Genomes of the genus *Tantillipelagibacter* occupy a previously uncharacterized monophyletic node within the Ia.4 cluster of the family *Pelagibacteraceae*, and share a minimum gANI of ~87% to the type genome. Cells of this genus have been isolated from the central Red Sea and tropical Pacific Ocean. The type species is *Tantillipelagibacter marisrubii.*

Registry URL: https://seqco.de/i:51401

**Description of *Proventipelagibacter* gen. nov.**

*Proventipelagibacter* ([Pro.ven.ti.pe.la.gi.bac'ter] L. masc. n. *proventus*, increase, yield, crop; L. neut. n. *pelagus*, the open sea; N.L. masc. n. *bacter*, a rod; N.L. masc. n. *Proventipelagibacter*, a marine bacterium with increased yield, referring to its increase in detection in select samples).

Genomes of the genus *Proventipelagibacter* occupy a previously uncharacterized monophyletic node within the Ia.4 cluster of the family *Pelagibacteraceae*, and share a minimum gANI of ~88% to the type genome. There are no known cultivated isolates yet reported for this genus. The type species is *Proventipelagibacter oceanicus.*

Registry URL: https://seqco.de/i:51403

**Description of *Punctipelagibacter* gen. nov.**

*Punctipelagibacter* ([Punc.ti.pe.la.gi.bac'ter] L. neut. n. *punctum*, a point, a spot; L. neut. n. *pelagus*, the open sea; N.L. masc. n. *bacter*, a rod; N.L. masc. n. *Punctipelagibacter*, a bacterium with spotty distribution globally).

Genomes of the genus *Punctumpelagibacter* occupy a previously uncharacterized monophyletic node within the Ia.4 cluster of the family *Pelagibacteraceae*, and share a minimum gANI of ~93% to the type genome. There are no known cultivated isolates yet reported for this genus. The type species is *Punctumpelagibacter variabilis.*

Registry URL: https://seqco.de/i:51405

**Description of *Pantolocipelagibacter* gen. nov.**

*Pantolocipelagibacter* ([Pan.to.lo.ci.pe.la.gi.bac'ter] Gr. masc. adj. *pas*, whole; L. masc. n. *locus*, a place; L. neut. n. *pelagus*, the open sea; N.L. masc. n. *bacter*, a rod; N.L. masc. n. *Pantolocipelagibacter*, a bacterium found across multiple latitudes and locations).

Genomes of the genus *Pantolocipelagibacter* occupy a previously uncharacterized monophyletic node within the Ia.4 cluster of the family *Pelagibacteraceae*, and share a minimum gANI of ~94% to the type genome. There are no known cultivated isolates yet reported for this genus. The type species *Pantolocipelagibacter undulae*.

Registry URL: https://seqco.de/i:51407

**Description of *Alipelagibacter* gen. nov.**

*Alipelagibacter* ([A.li.pe.la.gi.bac'ter] L. masc. pron. *alius*, other; L. neut. n. *pelagus*, the open sea; N.L. masc. n. *bacter*, a rod; N.L. masc. n. *Alipelagibacter*, the other bacterium from the open sea, referring to the presence of unique (or "other") molybdenum enzymes not found in closely related species).

Genomes of the genus *Alipelagibacter* occupy a previously uncharacterized monophyletic node within the Ia.4 cluster of the family *Pelagibacteraceae*. Its members encode the genetic pathway for synthesizing the molybdenum cofactor, while close evolutionary neighbors do not[16]. This genus is currently represented by a single isolate genome from the tropical Pacific Ocean. The type species is *Alipelagibacter loemeaikii.*

Registry URL: https://seqco.de/i:51409

**Description of *Lacunipelagibacter* gen. nov.**

*Lacunipelagibacter* ([La.cu.ni.pe.la.gi.bac'ter] L. fem. n. *lacuna*, a hole, an empty space, deficiency, loss; L. neut. n. *pelagus*, the open sea; N.L. masc. n. *bacter*, a rod; N.L. masc. n. *Lacunipelagibacter*, a bacterium missing enzymes that closely related groups harbor).

Genomes of the genus *Lacunipelagibacter* occupy a monophyletic node within the family *Pelagibacteraceae* that is currently known as SAR11 subclade Ia.4.II, and share a minimum gANI of ~90% with the type genome. Its members lack the genetic pathway required to synthesize the molybdenum cofactor, while some close evolutionary neighbors contain the required genes[16]. Cells of this genus have been isolated from the tropical Pacific Ocean. The type species is *Lacunipelagibacter aumakalihii.*

Registry URL: https://seqco.de/i:51411

**Description of *Coralipelagibacter* gen. nov.**

*Coralipelagibacter* ([Co.ra.li.pe.la.gi.bac'ter] L. neut. n. *coralium*, coral; L. neut. n. *pelagus*, the open sea; N.L. masc. n. *bacter*, a rod; N.L. masc. n. *Coralipelagibacter*, a bacterium that is coastally restricted where coral reefs are found).

Genomes of the genus *Coralipelagibacter* occupy a monophyletic node within the family *Pelagibacteraceae* that is currently known as SAR11 subclade Ia.3.II, and share a minimum gANI of ~84% to the type genome. Cells of this genus have been isolated from the tropical Pacific Ocean and this group is restricted globally to nearshore waters. The type species is *Coralipelagibacter heeiaensis.*

Registry URL: https://seqco.de/i:51413

**Description of *Litoralipelagibacter* gen. nov.**

*Litoralipelagibacter* ([Li.to.ra.li.pe.la.gi.bac'ter] L. masc. adj. *litoralis*, of the seashore; L. neut. n. *pelagus*, the open sea; N.L. masc. n. *bacter*, a rod; N.L. masc. n. *Litoralipelagibacter*, a coastal bacterium).

Genomes of the genus *Litoralipelagibacter* occupy a monophyletic node within the family *Pelagibacteraceae* that is currently known as SAR11 subclade Ia.3.III, and share a minimum gANI of ~95% to the type genome. Cells of this genus have been isolated from the tropical Pacific Ocean. The type species is *Litoralipelagibacter koolauensis.*

Registry URL: https://seqco.de/i:51416

**Description of *Stellipelagibacter* gen. nov.**

*Stellipelagibacter* ([Ste.lli.pe.la.gi.bac'ter] L. fem. n. *stella*, a star; L. neut. n. *pelagus*, the open sea; N.L. masc. n. *bacter*, a rod; N.L. masc. n. *Stellipelagibacter*, a bacterium of the starry sea, referring to the broad distribution of these bacteria as the vast starry sky).

Genomes of the genus *Stellipelagibacter* occupy a monophyletic node within the family *Pelagibacteraceae* that is currently known as SAR11 subclade Ib.2.I, and share a minimum gANI of ~85% to the type genome. There are no known cultivated isolates yet reported for this genus. The type species is *Stellipelagibacter aequoris.*

Registry URL: https://seqco.de/i:51419

**Description of *Parvipelagibacter* gen. nov.**

*Parvipelagibacter* ([Par.vi.pe.la.gi.bac'ter] L. masc. adj. *parvus*, small; L. neut. n. *pelagus*, the open sea; N.L. masc. n. *bacter*, a rod; N.L. masc. n. *Parvipelagibacter*, a bacterium with a tight and constrained distribution, which is smaller than that of its close neighbor.)

Genomes of the genus *Parvipelagibacter* occupy a previously uncharacterized monophyletic node within the family *Pelagibacteraceae*, and share a minimum gANI of ~90% to the type genome. There are no known cultivated isolates yet reported for this genus. The type species is *Parvipelagibacter dignotus.*

Registry URL: https://seqco.de/i:51421

**Description of *Superipelagibacter* gen. nov.**

*Superipelagibacter* ([Su.pe.ri.pe.la.gi.bac'ter] L. masc. adj. *superus*, above; L. neut. n. *pelagus*, the open sea; N.L. masc. n. *bacter*, a rod; N.L. masc. n. *Superipelagibacter*, a bacterium that is often correlated with higher temperatures).

Genomes of the genus *Superipelagibacter* occupy a monophyletic node within the family *Pelagibacteraceae* that is currently known as SAR11 subclade Ib.1.I, and share a minimum gANI of ~84% to the type genome. Cells of this genus have been isolated from the Red Sea and the tropical Pacific Ocean. The type species is *Superipelagibacter microcenae.*

Registry URL: https://seqco.de/i:51423

**Description of *Copipelagibacter* gen. nov.**

*Copipelagibacter* ([Co.pi.pe.la.gi.bac'ter] L. fem. n. *copia*, abundance, supply; L. neut. n. *pelagus*, the open sea; N.L. masc. n. *bacter*, a rod; N.L. masc. n. *Copipelagibacter*, an abundant rod from the open sea).

Genomes of the genus *Copipelagibacter* occupy a monophyletic node within the family *Pelagibacteraceae* that is currently known as SAR11 subclade Ib.1.III, and share a minimum gANI of ~90% to the type genome. There are no known cultivated isolates yet reported for this genus. The type species is *Copipelagibacter splendidus.*

Registry URL: https://seqco.de/i:51425

**Description of *Fusipelagibacter* gen. nov.**

*Fusipelagibacter* ([Fu.si.pe.la.gi.bac'ter] L. masc. adj. *fusus*, spread out; L. neut. n. *pelagus*, the open sea; N.L. masc. n. *bacter*, a rod; N.L. masc. n. *Fusipelagibacter*, a bacterium spread across the global oceans).

Genomes of the genus *Fusipelagibacter* occupy a previously uncharacterized monophyletic node within the Ib.4 genome cluster of the family *Pelagibacteraceae*, and share a minimum gANI of ~93% to the type genome. There are no known cultivated isolates yet reported for this genus. The type species is *Fusipelagibacter tranquillus.*

Registry URL: https://seqco.de/i:51427

**Description of *Extensipelagibacter* gen. nov.**

*Extensipelagibacter* ([Ex.ten.si.pe.la.gi.bac'ter] L. masc. perf. part. *extensus*, spread out; L. neut. n. *pelagus*, the open sea; N.L. masc. n. *bacter*, a rod; N.L. masc. n. *Extensipelagibacter*, a bacterium widely distributed across the ocean).

Genomes of the genus *Extensipelagibacter* occupy a monophyletic node within the family *Pelagibacteraceae* that is currently known as SAR11 subclade Ib.4.I, and share a minimum gANI of ~96% to the type genome. There are no known cultivated isolates yet reported for this genus. The type species is *Extensipelagibacter unicus.*

Registry URL: https://seqco.de/i:51429

**Description of *Medipelagibacter* gen. nov.**

*Medipelagibacter* ([Me.di.pe.la.gi.bac'ter] L. masc. adj. *medius*, middle; L. neut. n. *pelagus*, the open sea; N.L. masc. n. *bacter*, a rod; N.L. masc. n. *Medipelagibacter*, an open-ocean bacterium generally found at mid and low latitudes).

Genomes of the genus *Medipelagibacter* occupy a previously uncharacterized monophyletic node within the Ib.4 genome cluster of the family *Pelagibacteraceae*, and share a minimum gANI of ~84% to the type genome. There are no known cultivated isolates yet reported for this genus. The type species is *Medipelagibacter bermudensis.*

Registry URL: https://seqco.de/i:51431

**Description of *Amplipelagibacter* gen. nov.**

Amplipelagibacter ([Am.pli.pe.la.gi.bac'ter] L. masc. adj. *amplus*, great in number; L. neut. n. *pelagus*, the open sea; N.L. masc. n. *bacter*, a rod; N.L. masc. n. *Amplipelagibacter*, an abundant rod from the sea).

Genomes of the genus *Amplipelagibacter* occupy a previously uncharacterized monophyletic node within the Ib.4 genome cluster of the family *Pelagibacteraceae*, and share a minimum gANI of ~90% to the type genome. There are no known cultivated isolates yet reported for this genus. The type species is *Amplipelagibacter globalis.*

Registry URL: https://seqco.de/i:51433

**Description of *Rubripelagibacter* gen. nov.**

*Rubripelagibacter* ([Ru.bri.pe.la.gi.bac'ter] L. masc. adj. *ruber*, red, ruddy; L. neut. n. *pelagus*, the open sea; N.L. masc. n. *bacter*, a rod; N.L. masc. n. *Rubripelagibacter*, a bacterium of the red sea, referring to the Red Sea in which this genus is often found).

Genomes of the genus *Rubripelagibacter* occupy a previously uncharacterized monophyletic node within the family *Pelagibacteraceae*, and share a minimum gANI of ~85% to the type genome. There are no known cultivated isolates yet reported for this genus. The type species is *Rubripelagibacter caeruleus.*

Registry URL: https://seqco.de/i:51435

**Description of *Cosmipelagibacter* gen. nov.**

*Cosmipelagibacter* ([Cos.mi.pe.la.gi.bac'ter] L. masc. n. *cosmos*, universe; L. neut. n. *pelagus*, the open sea; N.L. masc. n. *bacter*, a rod; N.L. masc. n. *Cosmipelagibacter*, a bacterium found throughout the global oceans).

Genomes of the genus *Cosmipelagibacter* occupy a previously uncharacterized monophyletic node within the IIa.B genome cluster of the family *Cosmipelagibacteraceae*. A single strain from the tropical Pacific Ocean has been isolated to date. The type species is *Cosmipelagibacter regressus.*

Registry URL: https://seqco.de/i:51437

**Description of *Elongatipelagibacter* gen. nov.**

*Elongatipelagibacter* ([E.lon.ga.ti.pe.la.gi.bac'ter] L. masc. perf. part. *elongatus*, elongated; L. neut. n. *pelagus*, the open sea; N.L. masc. n. *bacter*, a rod; N.L. masc. n. *Elongatipelagibacter*, an elongated rod from the open sea, referring to the elongated morphology of this genus compared to other isolates from the *Pelagibacterales*).

Genomes of the genus *Elongatipelagibacter* occupy a previously uncharacterized monophyletic node within the IIIa.I genome cluster of the family *Fontibacteriaceae*. Multiple strains from this genus have been isolated. The type species is *Elongatipelagibacter longiformis.*

Registry URL: https://seqco.de/i:51439

**Description of *Xanthipelagibacter aumoanae* sp. nov.**

*Xanthipelagibacter aumoanae* ([a.u.mo.a'nae] N.L. gen. n. *aumoanae*, pertaining to one who swims the ocean, derived from the ʻŌlelo Hawaiʻi "au moana" meaning "swims the ocean").

The reference strain, HIMB83$^{T}$, was isolated from seawater taken from a depth of 2 m depth within Kāneʻohe Bay, Oʻahu, HI, USA at the Hawaiʻi Institute of Marine Biology located on the island of Moku o Loʻe, in the tropical Pacific Ocean. The DNA G+C content of the reference strain is 29.2 mol% (determined from the genome sequence), and the genome is 1.4 Mb in length. The GenBank accession number for the genome sequence of strain HIMB83$^{T}$ is GCA_000504225.1.

Registry URL: https://seqco.de/i:51393

**Description of *Undatipelagibacter huakaii* sp. nov.**

*Undatipelagibacter huakai* ([hu.a.ka'i.i] N.L. gen. n. *huakaii*, of a microbe from (water) traveling around coasts, from the ʻŌlelo Hawaiʻi "huakaʻi" meaning "traveling, around coasts").

The reference strain, HIMB1556$^{T}$, was isolated from seawater taken from a depth of 2 m within Kāneʻohe Bay, Oʻahu, HI, USA at the Hawaiʻi Institute of Marine Biology located on the island of Moku o Loʻe, in the tropical Pacific Ocean. The DNA G+C content of the reference strain is 29.3 mol% (determined from the genome sequence), and the genome is 1.5 Mb in length. The GenBank accession number for the genome sequence of strain HIMB1556$^{T}$ is JBNAJE000000000.

Registry URL: https://seqco.de/i:51395

**Description of *Angustipelagibacter atlanticus* sp. nov.**

*Angustipelagibacter atlanticus* ([a.tlan'ti.cus] L. masc. adj. *atlanticus*, Atlantic, from the Atlantic ocean).

The type genome, SAG GCA_902580005.1$^{T}$, was collected from seawater taken from a depth of 89.9 m within the Mid-Atlantic Ocean. The DNA G+C content of the type genome is 29.8 mol% (determined from the genome sequence), and the genome is 1.18 Mb in length. The GenBank accession number for the genome sequence is GCA_902580005.1.

Registry URL: https://seqco.de/i:51397

**Description of *Atlantikopelagibacter sargassoseensis* sp. nov.**

*Atlantikopelagibacter sargassoseensis* ([sar.gas.so.se.en'sis] N.L. masc. adj. *sargassoseensis*, of the Sargasso Sea, the site from which multiple strains of this species have been isolated).

The reference strain, HTCC7211$^{T}$, was isolated from seawater taken from a depth of 10 m within the western Atlantic Ocean. The DNA G+C content of the reference strain is 29.0 mol% (determined from the genome sequence), and the genome is 1.5 Mb in length. The

GenBank accession number for the genome sequence of strain HTCC7211$^T$ is GCF_000155895.1.

Registry URL: https://seqco.de/i:49682

**Description of *Accedentipelagibacter yaqonii* sp. nov.**

*Accedentipelagibacter yaqonii* ([ya.qo'ni.i] N.L. gen. n. *yaqonii*, of the Yaqo'n (pronounced Yacona) people that lived along the coast of the Northeastern Pacific Ocean in what is now the central coast of Oregon, USA).

The reference strain, HTCC8051$^T$, was isolated from seawater taken from a depth of 10 m off of the coast of Newport, OR, USA, in the northeast Pacific Ocean. The DNA G+C content of the reference strain is 29.2 mol% (determined from the genome sequence), and the genome is 1.4 Mb in length. The GenBank accession number for the genome sequence of strain HTCC8051$^T$ is GCA_000472605.1.

Registry URL: https://seqco.de/i:51399

**Description of *Colligatipelagibacter wilhelmii* sp. nov.**

*Colligatipelagibacter wilhelmii* ([wil.hel'mi.i] N.L. masc. gen. n. *wilhelmii*, of Wilhelm, named after Dr. Larry Wilhelm, instrumental in early SAR11 research).

The type genome, SAG GCA_902524535.1$^T$, was collected from seawater taken from a depth of 10 m in the western Atlantic Ocean. The DNA G+C content of the type genome is 29.4 mol% (determined from the genome sequence), and the genome is 1.1 Mb in length. The GenBank accession number for the genome sequence is GCA_902524535.1.

Registry URL: https://seqco.de/i:49684

**Description of *Tantilluspelagibacter marisrubii* sp. nov.**

*Tantilluspelagibacter marisrubii* ([ma.ris.ru'bi.i] L. neut. n. *mare*, the sea; L. masc. adj. *rubius*, red; N.L. gen. n. *marisrubii*, of the Red Sea, where the first strain was isolated from).

The reference strain, HIMB1436$^T$, was isolated from seawater taken from a depth of 2 m depth from outside of Kāneʻohe Bay, Oʻahu, HI, USA, at the Hawaiʻi Institute of Marine Biology located on the island of Moku o Loʻe, in the tropical Pacific Ocean.. While the first isolated strain (RS39) originated from the central Red Sea at 19 m depth, it was not eligible to be submitted to the SeqCode as a reference strain as the raw DNA genome sequence reads are no longer available. The DNA G+C content of the reference strain is 29.2 mol% (determined from the genome sequence), and the genome is 1.2 Mb in length. The GenBank accession number for the genome sequence of strain HIMB1436$^T$ is JBNGZC000000000.

Registry URL: https://seqco.de/i:51402

**Description of *Proventipelagibacter oceanicus* sp. nov.**

*Proventipelagibacter oceanicus* ([o.ce.a'ni.cus] N.L. masc. adj. *oceanicus*, of the ocean).

The type genome, SAG GCA_902621345.1$^T$, was collected from seawater taken from a depth of 10 m within the western Atlantic Ocean. The DNA G+C content of the type genome is 29.5 mol% (determined from the genome sequence), and the genome is 1.2 Mb in length. The GenBank accession number for the genome sequence is GCA_902621345.1.

Registry URL: https://seqco.de/i:51404

**Description of *Punctumpelagibacter variabilis* sp. nov.**

*Punctumpelagibacter variabilis* ([va.ri.a.bi'lis] L. masc. adj. *variabilis*, variable, referring to a variable global distribution).

The type genome, SAG GCA_902540275.1$^{T}$, was collected from seawater taken from a depth of 10 m within the western Atlantic Ocean. The DNA G+C content of the genome is 28.3 mol% (determined from the genome sequence), and the genome is 1.2 Mb in length. The GenBank accession number for the genome sequence is GCA_902540275.1.

Registry URL: https://seqco.de/i:51406

**Description of *Pantolocipelagibacter undulae* sp. nov.**

*Pantolocipelagibacter undulae* ([un.du'lae] L. gen. n. *undulae*, of small waves, referring to the marine habitat of this bacterium).

The type genome, SAG GCA_902597885.1$^{T}$, was collected from seawater taken from a depth of 10 m within the western Atlantic Ocean. The DNA G+C content of the type genome is 28.8 mol% (determined from the genome sequence), and the genome is 1.1 Mb in length. The GenBank accession number for the genome sequence is GCA_902597885.1.

Registry URL: https://seqco.de/i:51408

**Description of *Alipelagibacter loemeaiki* sp. nov.**

*Alipelagibacter loemeaikii* ([lo.e.me.a.i'ki.i] N.L. gen. n. *loemeaikii*, of a small organism of Lo'e, derived from the ʻŌlelo Hawaiʻi "Lo'e mea iki" meaning "a small organism from Lo'e", referring to the island of Moku o Loʻe).

The reference strain, HIMB1483$^{T}$, was isolated from seawater taken from a depth of 2 m within Kāneʻohe Bay, Oʻahu, HI, USA, at the Hawaiʻi Institute of Marine Biology located on the island of Moku o Loʻe, in the tropical Pacific Ocean. The DNA G+C content of the reference strain is 28.5 mol% (determined from the genome sequence), and the genome is 1.2 Mb in length. The GenBank accession number for the genome sequence of strain HIMB1483$^{T}$ is JBNAIM000000000.

Registry URL: https://seqco.de/i:51410

**Description of *Lacunipelagibacter aumakalihii* sp. nov.**

*Lacunipelagibacter aumakalihii* ([a.u.ma.ka.li'hi.i] N.L. gen. n. *aumakalihii*, of a microbe swimming the outside border, from the ʻŌlelo Hawaiʻi "au ma ka lihi" meaning "swims the outside border").

The reference strain, HIMB1437$^{T}$, was isolated from seawater taken from a depth of 2 m within Kāneʻohe Bay, Oʻahu, HI, USA at the Hawaiʻi Institute of Marine Biology located on the island of Moku o Loʻe, in the tropical Pacific Ocean. The DNA G+C content of the reference strain is 29.4 mol% (determined from the genome sequence), and the genome is 1.2 Mb in length. The GenBank accession number for the genome sequence of strain HIMB1437$^{T}$ is JBNGZD000000000.

Registry URL: https://seqco.de/i:51412

**Description of *Coralipelagibacter heeiaensis* sp. nov.**

*Coralipelagibacter heeiaensis* ([he.e.i.a.en'sis] N.L. masc. adj. *heeiaensis*, isolated from the ahupuaʻa (a traditional subdivision of land and sea) of Heʻeia off the island of Oʻahu).

The reference strain, HIMB5$^{T}$, was isolated from seawater taken from a depth of 2 m within Kāneʻohe Bay, Oʻahu, HI, USA, at the Hawaiʻi Institute of Marine Biology located on the island of Moku o Lo‘e, in the tropical Pacific Ocean. The DNA G+C content of the reference strain is 28.6 mol% (determined from the genome sequence), and the genome is 1.3 Mb in length. The GenBank accession number for the genome sequence of strain HIMB5$^T$ is GCF_000299095.1.

Registry URL: https://seqco.de/i:51414

**Description of *Coralipelagibacter kukahii* sp. nov.**

*Coralipelagibacter kukahii* ([k.uka'hi.i] N.L. gen. n. *kukahii*, referring to a state of uniqueness, derived from the ‘Ōlelo Hawai‘i "kū kahi" meaning "unique, standing alone").

The reference strain, HIMB1412$^T$, was isolated from seawater taken from a depth of 2 m within Kāne‘ohe Bay, O‘ahu, HI, USA, at the Hawai‘i Institute of Marine Biology located on Moku o Lo‘e, in the tropical Pacific Ocean. The DNA G+C content of the reference strain is 29.3 mol% (determined from the genome sequence), and the genome is 1.3 Mb in length. The GenBank accession number for the genome sequence of strain HIMB1412$^T$ is JBNAKG000000000.

Registry URL: https://seqco.de/i:51415

**Description of *Litoralipelagibacter koolauensis* sp. nov.**

*Litoralipelagibacter koolauensis* ([ko.o.la.u.en'sis] N.L. masc. adj. *koolauensis*, of Ko‘olau, which is the ‘Ōlelo Hawai‘i term for "windward", referring in this case specifically both to Ko‘olau, the ancient eastern shield volcano on O‘ahu, and to the Ko‘olau Range, the modern-day remnants of the Ko‘olau volcano that overlook the bay from which the first isolate of this species was obtained).

The reference strain, HIMB4$^T$, was isolated from seawater taken from a depth of 2 m water depth within Kāne‘ohe Bay, O‘ahu, HI, USA at the Hawai‘i Institute of Marine Biology located on the island of Moku o Lo‘e, in the tropical Pacific Ocean. The DNA G+C content of the reference strain is 29.0 mol% (determined from the genome sequence), and the genome is 1.4 Mb in length. The GenBank accession number for the genome sequence of strain HIMB4$^T$ is JBPPFS000000000.

Registry URL: https://seqco.de/i:51417

**Description of *Pelagibacter yakuwasensis* sp. nov.**

*Pelagibacter yakuwasiensis* ([ya.ku.wa.si.en'sis] N.L. masc. adj. *yakuwasiensis*, of the vast ocean, derived from "yaku wasi" in the language of the Yaqo'n (pronounced Yacona) people that lived along the coast of the Northeastern Pacific Ocean in what is now the central coast of Oregon, USA).

The reference strain, HTCC9565$^T$, was isolated from surface seawater of the Pacific Ocean from over the Juan de Fuca Ridge off the coast of Newport, OR, USA. The DNA G+C content of the reference strain is 28.9 mol% (determined from the genome sequence), and the genome is 1.3 Mb in length. The GenBank accession number for the genome sequence of strain HTCC9565$^T$ is GCF_012932795.1.

Registry URL: https://seqco.de/i:51418

**Description of *Stellipelagibacter aequoris* sp. nov.**

*Stellipelagibacter aequoris* ([ae'quor.is] L. gen. n. *aequoris*, of the sea, referring to the broad distribution of this species).

The type genome, SAG GCA_902583075.1$^{T}$, was collected from seawater of the Atlantic Ocean at 10 m water depth. The DNA G+C content of the type genome is 29.4 mol% (determined from the genome sequence), and the genome is 1.2 Mb in length. The GenBank accession number for the genome sequence is GCA_902583075.1. This name refers to the broad distribution of this species.

Registry URL: https://seqco.de/i:51420

**Description of *Parvipelagibacter dignotus* sp. nov.**

*Parvipelagibacter dignotus* ([dig.no'tus] L. masc. part. adj. *dignotus*, dignified, distinguished, recognized as different from its closest neighbor).

The type genome, SAG GCA_902577075.1$^{T}$, was collected from seawater taken from a depth of 90.8 m within the northwest Atlantic Ocean. The DNA G+C content of the type genome is 29.4 mol% (determined from the genome sequence), and the genome is 1.36 Mb in length. The GenBank accession number for the genome sequence is GCA_902577075.1.

Registry URL: https://seqco.de/i:51422

**Description of *Superipelagibacter microcenae* sp. nov.**

*Superipelagibacter microcenae* ([mi.cro.ce'nae] Gr. masc. adj. *mikros*, small; L. fem. n. cena, meal; N.L. gen. n. *microcenae*, of the small meal, referring to the utilization of small compounds including aldehydes and formate)[16].

The reference strain, HIMB2305$^{T}$, was isolated from seawater taken from a depth of 2 m outside of Kāne‘ohe Bay, O‘ahu, HI, USA at the Hawai‘i Institute of Marine Biology located on the island of Moku o Lo‘e, in the tropical Pacific Ocean. While the first cultured isolate of this genus originated from the central Red Sea at 19 m water depth (RS40), it cannot serve as type material for the SeqCode as the raw reads are no longer available. The DNA G+C content of the reference strain HIMB2305$^{T}$ is 29.5 mol% (determined from the genome sequence), and the genome is 1.4 Mb in length. The GenBank accession number for the genome sequence of strain HIMB2305$^{T}$ is JBNAKJ000000000.

Registry URL: https://seqco.de/i:51424

**Description of *Copipelagibacter splendidus* sp. nov.**

*Copipelagibacter splendidus* ([splen.di'dus] L. masc. adj. *splendidus*, splendid, glittering).

The type genome, SAG GCA_902617415.1$^{T}$, was collected from seawater taken from a depth of 10 m in the western Atlantic Ocean. The DNA G+C content of the type genome is 29.2 mol% (determined from the genome sequence), and the genome is 1.4 Mb in length. The GenBank accession number for the genome sequence is GCA_902617415.1.

Registry URL https://seqco.de/i:51426

**Description of *Fusipelagibacter tranquillus* sp. nov.**

*Fusipelagibacter tranquillus* ([tran.qui'llus] L. masc. adj. *tranquillus*, quiet, calm).

The type genome, SAG GCA_902560075.1$^{T}$, was collected from seawater taken from a depth of 10 m within the western Atlantic Ocean. The DNA G+C content of the type genome is 29.1 mol% (determined from the genome sequence), and the genome is 1.2 Mb in length. The GenBank accession number for the genome sequence is GCA_902560075.1.

Registry URL: https://seqco.de/i:51428

**Description of *Extensipelagibacter unicus* sp. nov.**

*Extensipelagibacter unicus* ([u.ni'cus] L. masc. adj. *unicus*, unique; referring to the high genome similarity found between members of this species, a unique trait compared to closely related lineages).

The type genome, SAG GCA_902576855.1$^{T}$, was collected from seawater taken from a depth of 90.8 m within the northwest Atlantic Ocean. The DNA G+C content of the type genome is 28.7 mol% (determined from the genome sequence), and the genome is 1.2 Mb in length. The GenBank accession number for the genome sequence is GCA_902576855.1.

Registry URL: https://seqco.de/i:51430

**Description of *Medipelagibacter bermudensis* sp. nov.**

*Medipelagibacter bermudensis* ([ber.mu.den'sis] N.L. masc. adj. *bermudensis*, of Bermuda, referring to the sampling location near Bermuda at the Bermuda Atlantic Time-series (BATS) site).

The type genome, SAG GCA_902622545.1$^{T}$, was collected from seawater taken from a depth of 10 m within the western Atlantic Ocean. The DNA G+C content of the type genome is 28.9 mol% (determined from the genome sequence), and the genome is 1.2 Mb in length. The GenBank accession number for the genome sequence is GCA_902622545.1.

Registry URL: https://seqco.de/i:51432

**Description of *Amplipelagibacter globalis* sp. nov.**

*Amplipelagibacter globalis* ([glo.ba'lis] N.L. masc. adj. *globalis*, global, referring to the global distribution of the species).

The type genome, SAG GCA_902580675.1$^{T}$, was collected from seawater taken from a depth of 89.9 m within the Atlantic Ocean. The DNA G+C content of the type genome is 29.5 mol% (determined from the genome sequence), and the genome is 1.1 Mb in length. The GenBank accession number for the genome sequence is GCA_902580675.1.

Registry URL: https://seqco.de/i:51434

**Description of *Rubripelagibacter caeruleus* sp. nov.**

*Rubripelagibacter caeruleus* ([cae.ru.le'us] L. masc. adj. *caeruleus*, blue, cerulean, referring to the ocean environment).

The type genome, SAG GCA_902581445.1$^{T}$, was collected from seawater taken from a depth of 10 m within the western Atlantic Ocean. The DNA G+C content of the type genome is 29.2 mol% (determined from the genome sequence), and the genome is 1.1 Mb in length. The GenBank accession number for the genome sequence is GCA_902581445.1.

Registry URL: https://seqco.de/i:51436

**Description of *Cosmipelagibacter regressus* sp. nov.**

*Cosmipelagibacter regressus* ([re.gres'sus] L. gen. n. *regressus*, of going back, of the return, referring to the *Cosmiplelagibacter* that has returned (to the coast)).

The reference strain, HIMB58$^{T}$, was isolated from seawater taken from a depth of 2 m within Kāne‘ohe Bay, O‘ahu, HI, USA, at the Hawai‘i Institute of Marine Biology located on the island of Moku o Lo‘e, in the tropical Pacific Ocean. The DNA G+C content of the reference strain is 29.9 mol% (determined from the genome sequence), and the genome is 1.1 Mb in length. The GenBank accession number for the genome sequence of strain HIMB58$^T$ is GCA_000419545.1.

Registry URL: https://seqco.de/i:51438

**Description of *Elongatipelagibacter longiformis* sp. nov.**

*Elongatipelagibacter longiformis* ([lon.gi.for'mis] L. masc. adj. *longus*, long; L. masc. adj. suff. -*formis*, in the shape of; N.L. masc. adj. *longiformis*, long-shaped (compared to other *Pelagibacterales*)).

The reference strain, HIMB114$^T$, was isolated from seawater taken from a depth of 2 m within Kāneʻohe Bay, Oʻahu, HI, USA, at the Hawaiʻi Institute of Marine Biology located on the island of Moku o Loʻe in the tropical Pacific Ocean. The DNA G+C content of the reference strain is 29.6 mol% (determined from the genome sequence), and the genome is 1.2 Mb in length. The GenBank accession number for the genome sequence of strain HIMB114$^T$ is GCA_000163555.2.

Registry URL: https://seqco.de/i:51440

**Description of *Mesopelagibacter profundi* sp. nov.**

*Mesopelagibacter profundi* ([pro.fun'di] L. gen. n. *profundi*, of the depths, living within the depths of the oceans).

The type genome, SAG GCA_000504625.1$^T$, was collected from seawater taken from a depth of 770 m within the tropical Pacific Ocean from Station ALOHA (located at 22° 45'N, 158°W). The DNA G+C content of the type genome is 29.0 mol% (determined from the genome sequence), and the genome is 1.4 Mb in length.

Registry URL: https://seqco.de/i:52920

### Supplementary Figures

Supplementary Figures are available on FigShare at the following DOI: 10.6084/m9.figshare.30048979.

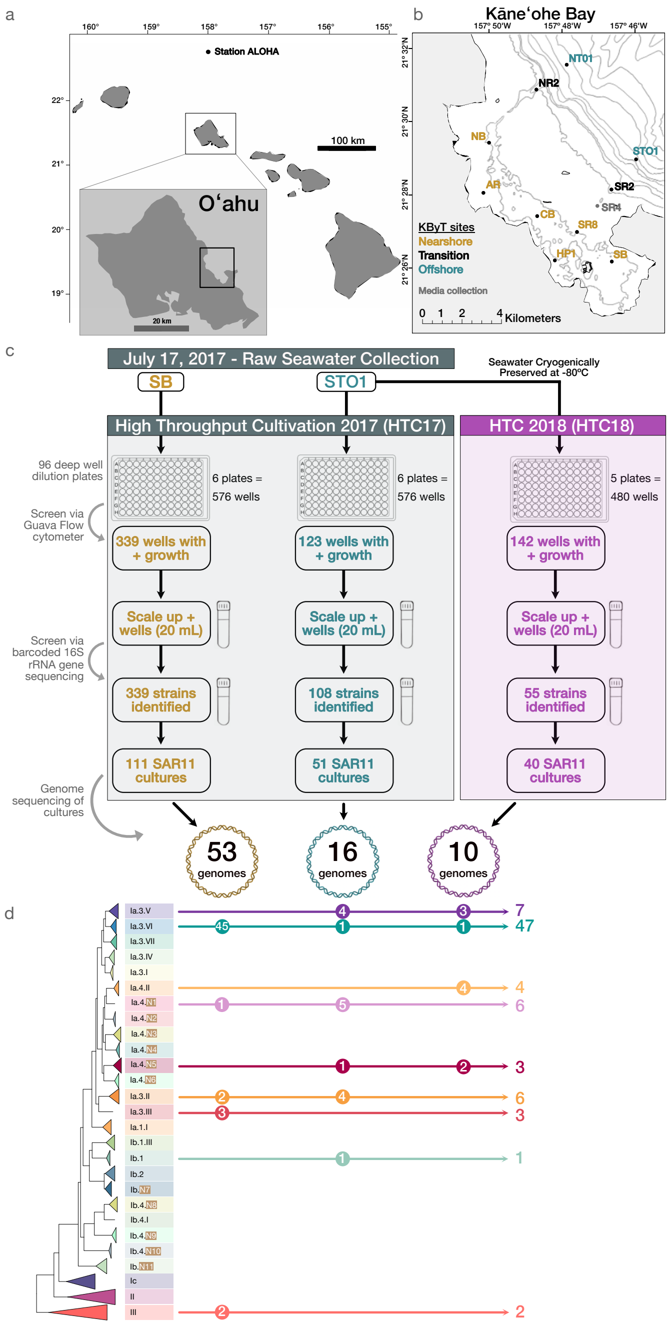


**Supplementary Fig. 1**. **Overview of high-throughput culturing experiments.** (**A**) Location of O‘ahu in the Hawaiian archipelago in relation to Station ALOHA (~100 km north). (**B**) Map of the embayment on the windward side of O‘ahu with sites included in the Kāne‘ohe Bay Time-series (KByT) that are classified as ‘nearshore’ (orange), ‘transition’ (black), or ‘offshore’ (turquoise). Site SR4 in gray from which seawater media was collected for the cultivation experiments is also indicated. Bathymetry lines are approximate indicating every 10 meters up to 50 meters, then marking 100 meter increments, after Tucker et al.,[28]. (**C**) Flowchart outlining the high throughput cultivation (HTC) experiments conducted in 2017 and 2018 leading to the isolation of hundreds of SAR11 cultures and 79 new SAR11 isolate genomes. (**C**) Schematic SAR11 phylogeny indicating which samples harbored genomes from which subgroups.

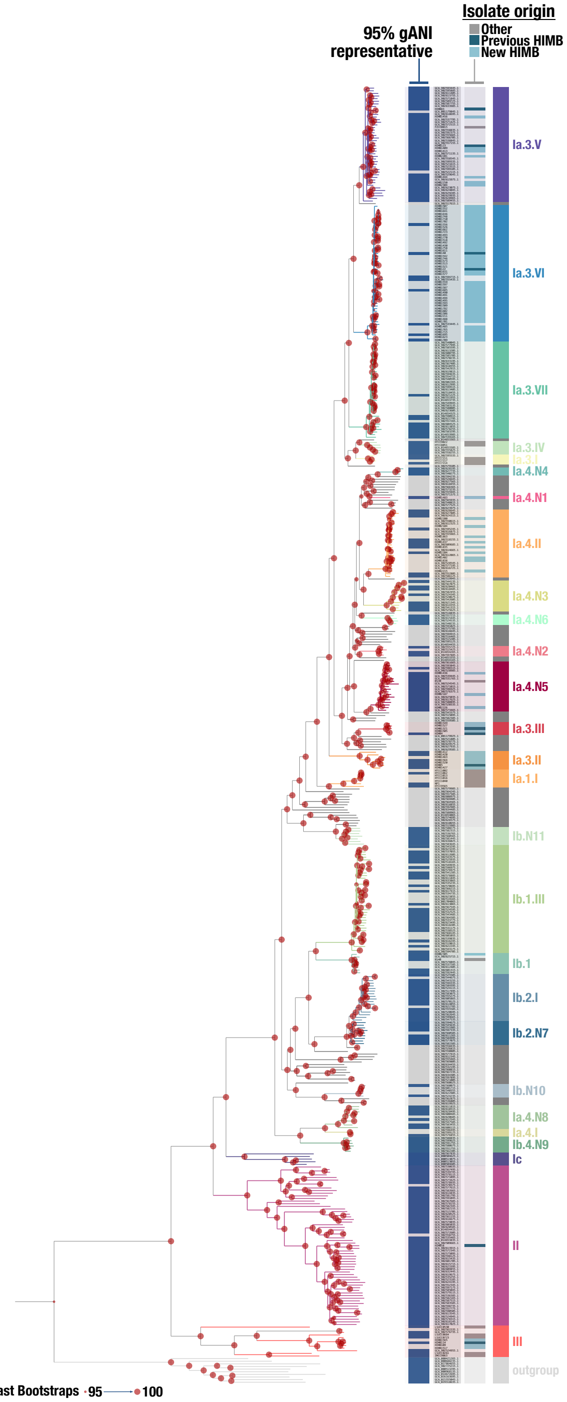


**Supplementary Fig. 2. Phylogenomic tree of all 484 *Pelagibacterales* genomes included with accession numbers.** Of the 484 SAR11 genomes 109 were isolates and 375 SAGs. The phylogeny is based on protein sequences from a curated SAR11-specific set of 165 genes. Isolate origin indicates if the genome was from this study, a previous isolate from Kāne‘ohe Bay, or from another source. Genomes that serve as a 95% gANI representative and were included in the read recruitment are also indicated with a dark blue bar. Circles indicate ultrafast bootstrap support values ≥95%, from 1000 replicates.

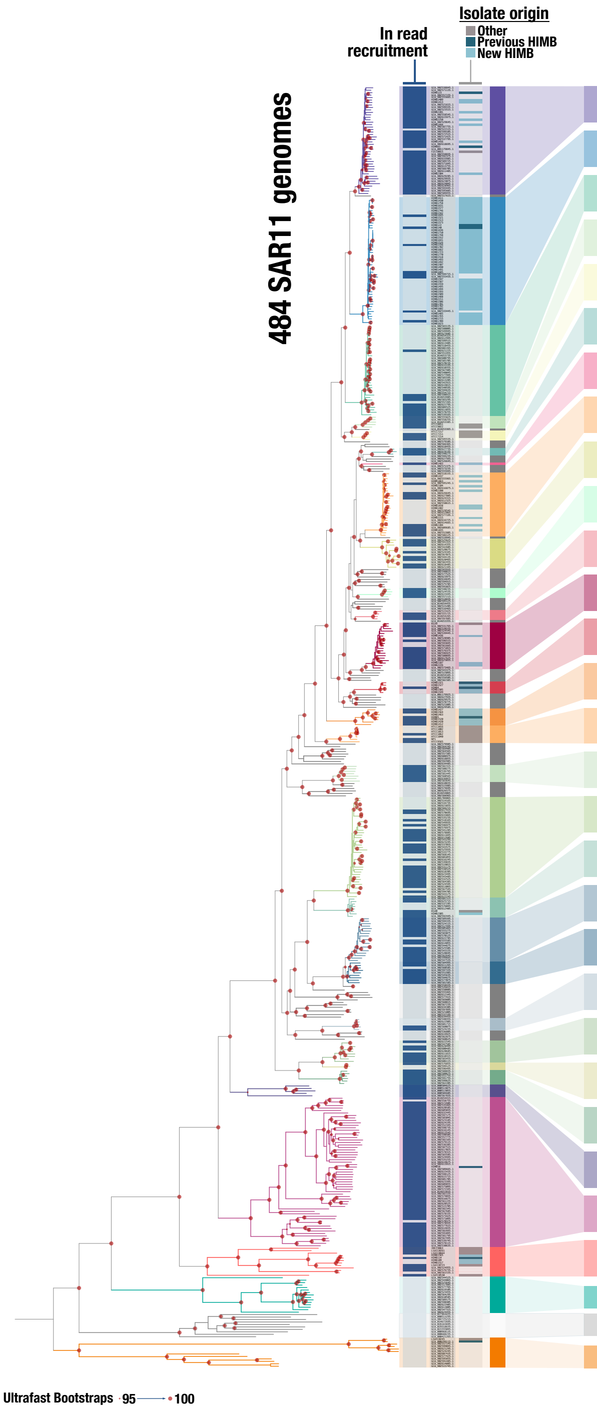


| | gANI (%) | | |
|---|---|---|---|
| | Min | Avg | Max |
| Ia.3.V | 84.9 | 89.8 | 95.9 |
| Ia.3.VI | 88.1 | 94.4 | 99.9 |
| Ia.3.VII | 85.7 | 93.6 | 99.9 |
| Ia.3.IV | 86.3 | 88.2 | 93.8 |
| Ia.3.I | 92.1 | 93.2 | 95.1 |
| Ia.4.N4 | 92.9 | 94.6 | 99.9 |
| Ia.4.N1 | - | - | - |
| Ia.4.II | 90.3 | 93.7 | 99.9 |
| Ia.4.N3 | 88.1 | 92.2 | 99.9 |
| Ia.4.N6 | 91.9 | 92.4 | 93.0 |
| Ia.4.N2 | 94.1 | 94.6 | 95.1 |
| Ia.4.N5 | 86.5 | 91.3 | 98.8 |
| Ia.3.III | 95.0 | 95.8 | 99.9 |
| Ia.3.II | 84.4 | 90.1 | 95.4 |
| Ia.1.I | 84.2 | 92.1 | 98.5 |
| Ib.N11 | 93.2 | 94.7 | 96.8 |
| Ib.1.III | 89.7 | 93.3 | 99.9 |
| Ib.1 | 90.3 | 93.0 | 96.0 |
| Ib.2.I | 85.2 | 89.5 | 99.9 |
| Ib.N7 | 89.5 | 91.4 | 99.9 |
| Ib.N10 | 90.1 | 91.6 | 92.6 |
| Ib.4.N8 | 83.7 | 90.0 | 99.9 |
| Ib.4.I | 96.4 | 96.9 | 97.4 |
| Ib.4.N9 | 90.1 | 91.6 | 92.6 |
| Ic | 77.3 | 82.6 | 86.9 |
| II | 76.2 | 80.1 | 99.9 |
| III | 76.6 | 86.0 | 96.2 |
| IV | | | |
| Rhodobacteraceae | | | |
| AEGEAN-169 (V) | | | |

**Supplementary Fig. 3. Phylogenomic tree with 484 *Pelagibacterales* genomes and subgroups IV and V genomes.** Isolate origin is highlighted on the tree and indicates if the genome was from this study, a previous isolate from Kāne'ohe Bay, or from another source. Genomes that serve as a 95% gANI representative and were included in the read recruitment are also indicated with a dark blue bar.

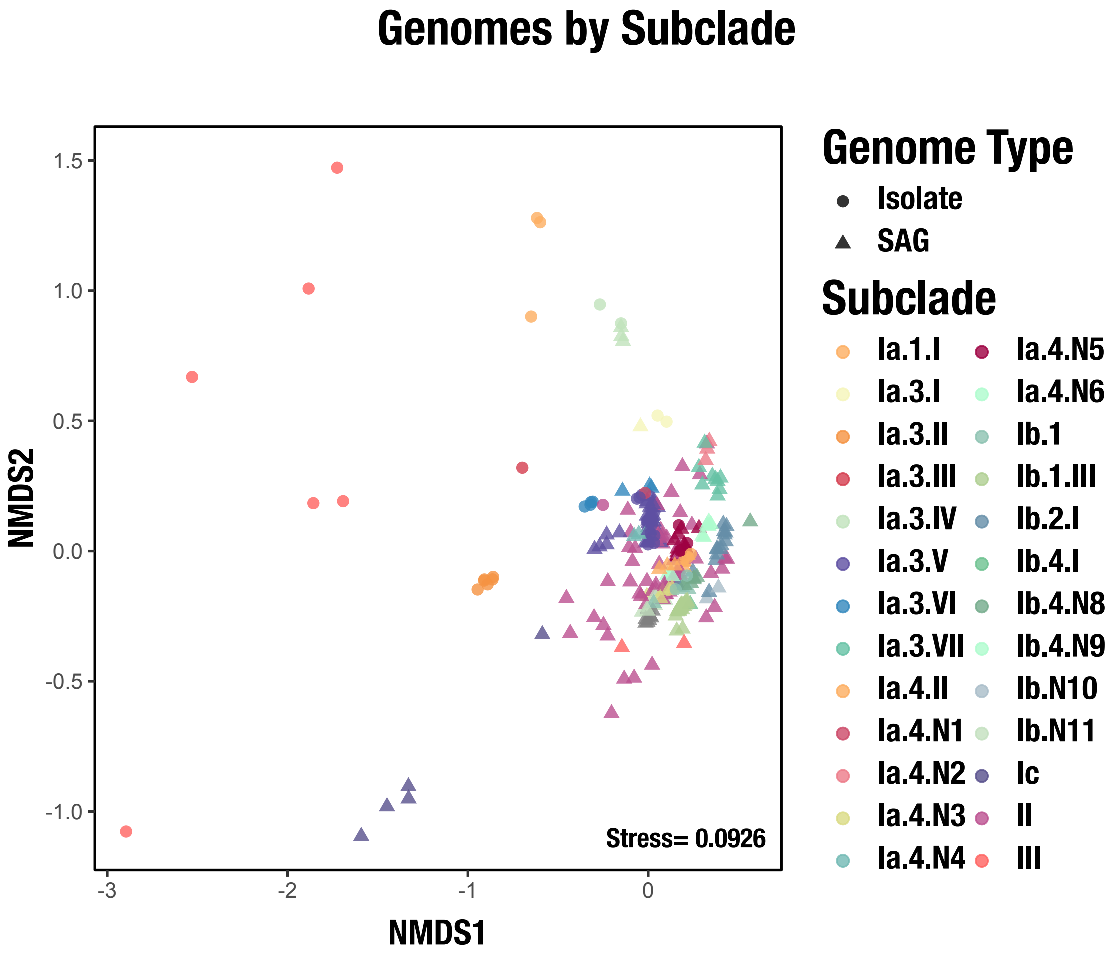


**Supplementary Fig. 4. NMDS of genomes with detection data.** All of the genomes included in the analysis are included here, with distinction between isolate genome (circle) or SAG (triangle) indicated.

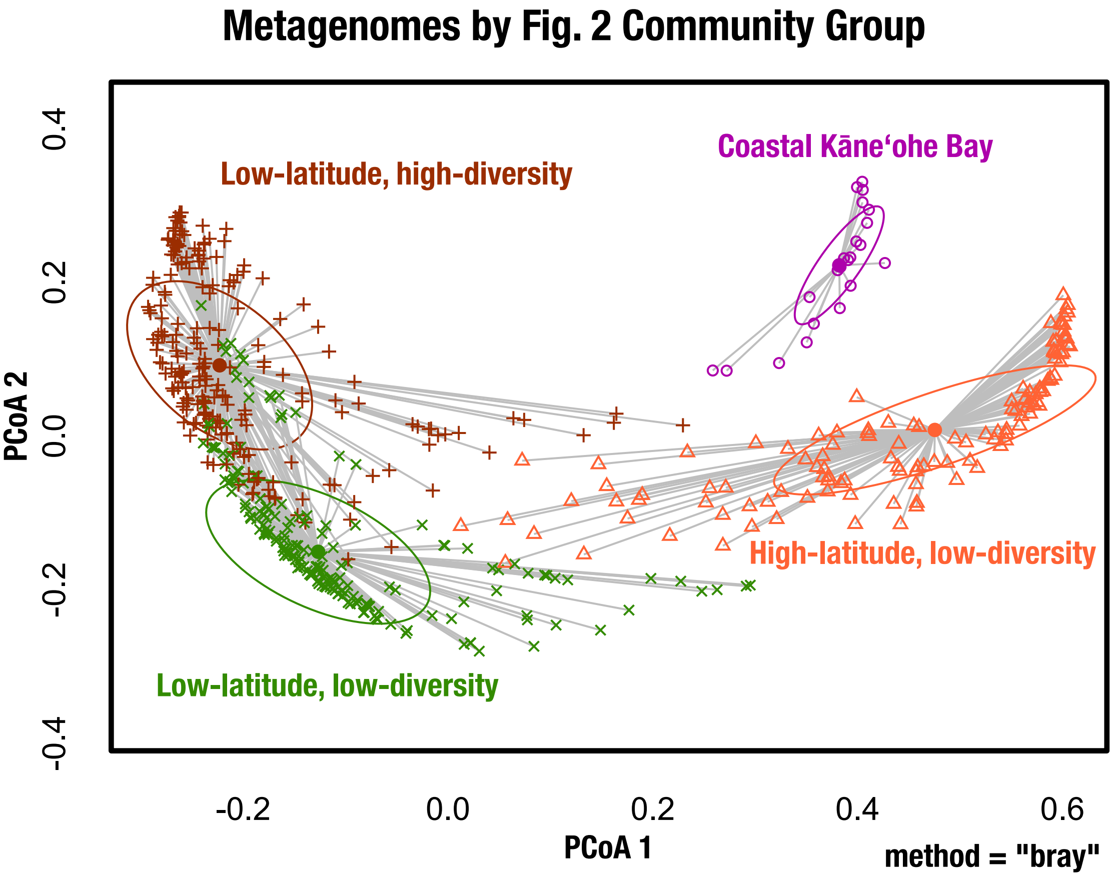


**Supplementary Fig. 5. PCoA of metagenomes by community group.** All of the genomes included in the read recruitment analysis are included here as well as all metagenomes in Figure 2. This is a visualization of the PERMDISP results.

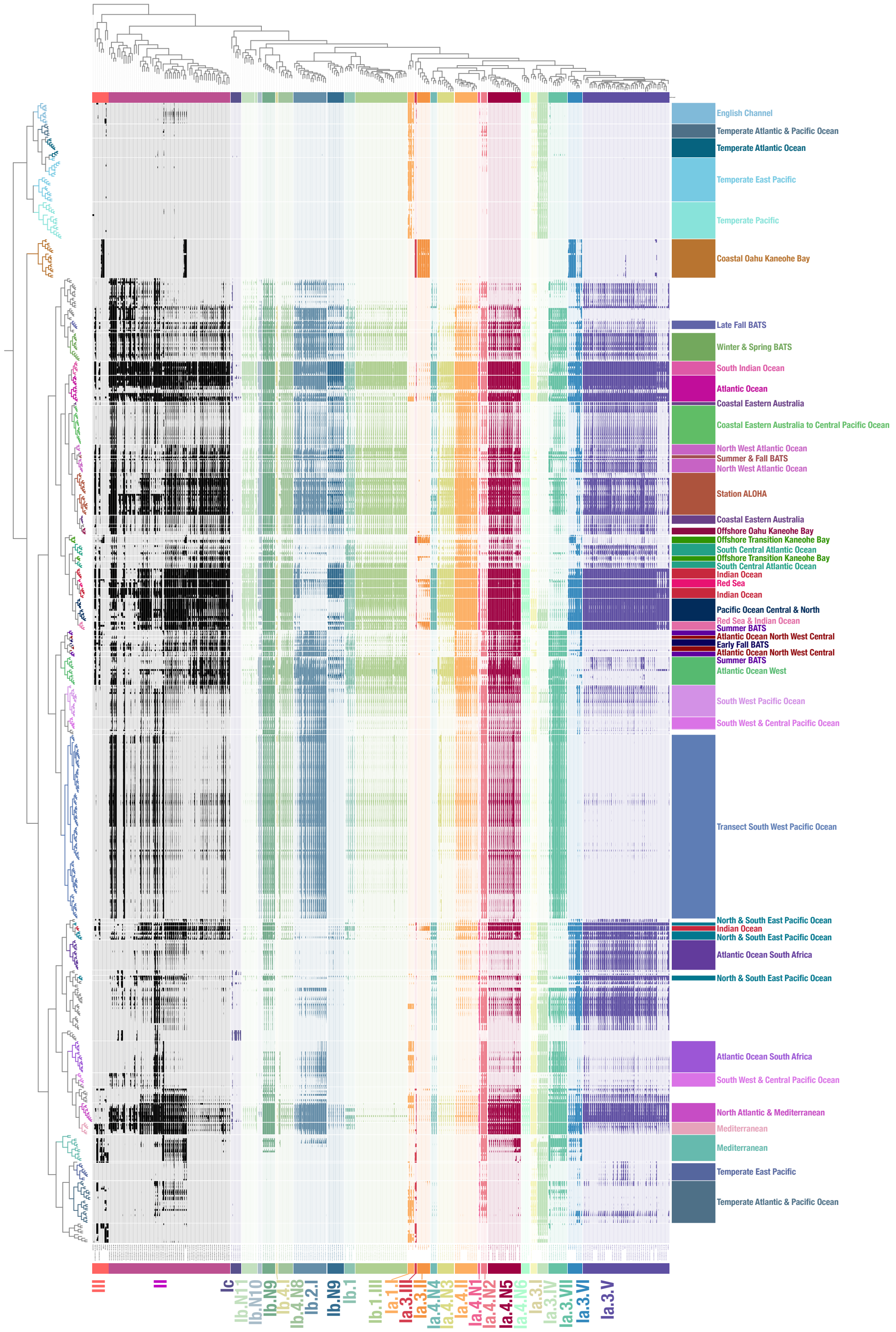


**Supplementary Fig. 6 Read recruitment data including all metagenomes included in cluster analysis of metagenomes based on genome detection values using the k-means algorithm.** Subgroups are indicated at the bottom of the figure with labels indicating the metagenome groups along the right hand side.

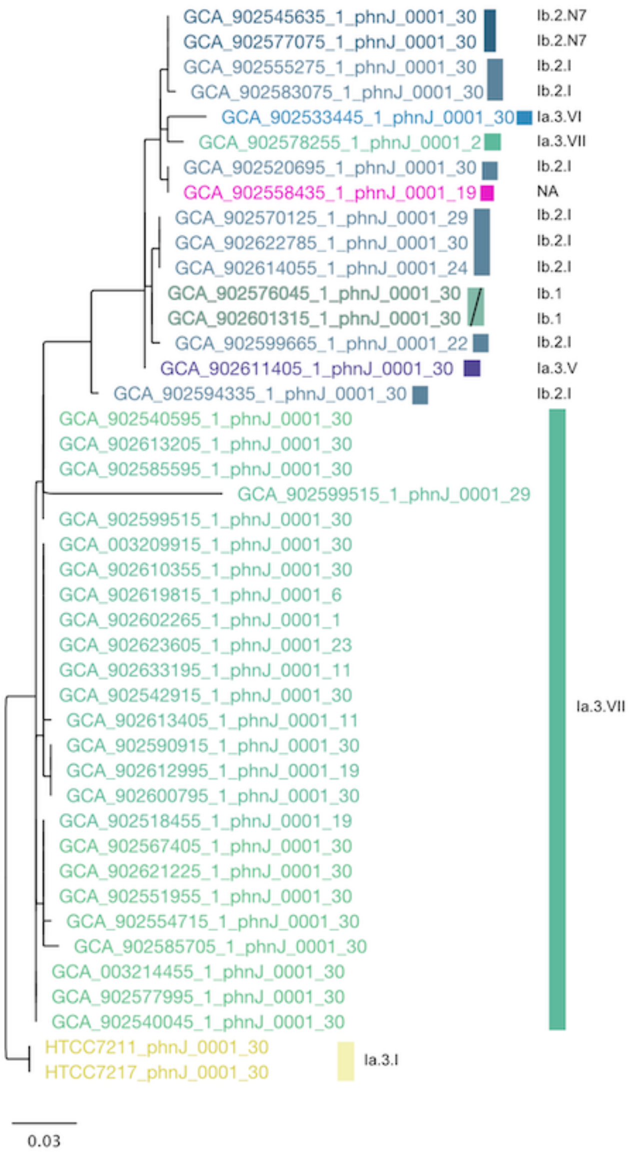


**Supplementary Fig. 7. Phylogeny of the *phnJ* gene.** A phylogeny of protein sequences from all *Pelagibacterales* genomes in the pruned phylogeny data set that contain the *phnJ* gene.

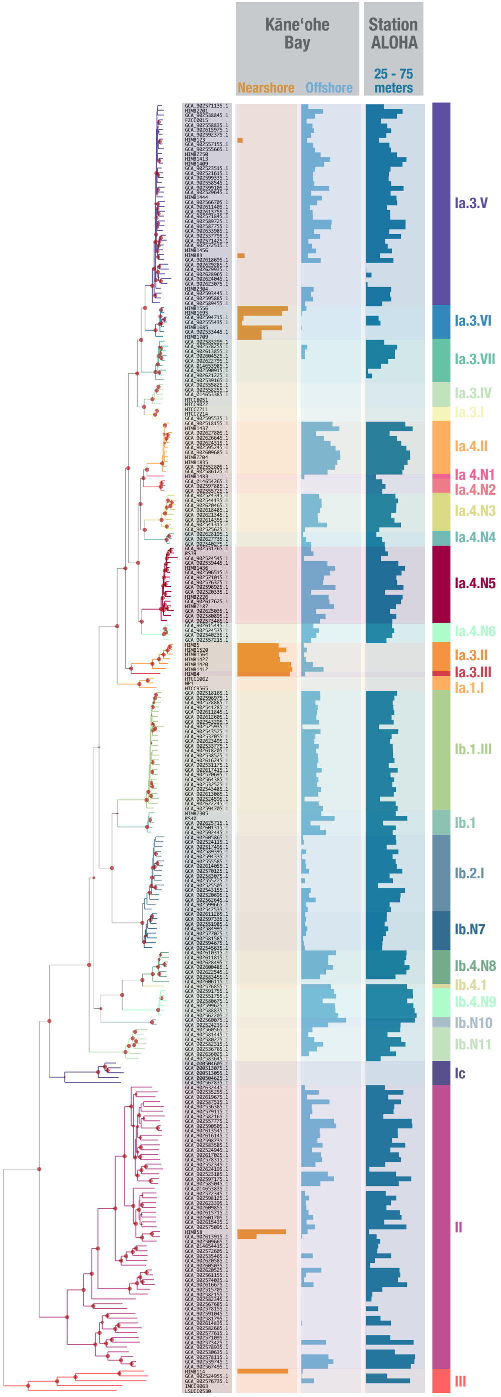


**Supplementary Fig. 8. Phylogenomic tree of the *Pelagibacterales* with read recruitment from Kāne'ohe Bay and Station ALOHA.** The phylogeny includes 268 genomes (50 isolates and 218 SAGs) included in the pruned phylogeny. Read recruitment data in the form of detection values on a scale of 0.25 to 0.75 are displayed along the right hand side. Circles indicate ultrafast bootstrap support values ≥95% or greater from 1000 replicates.

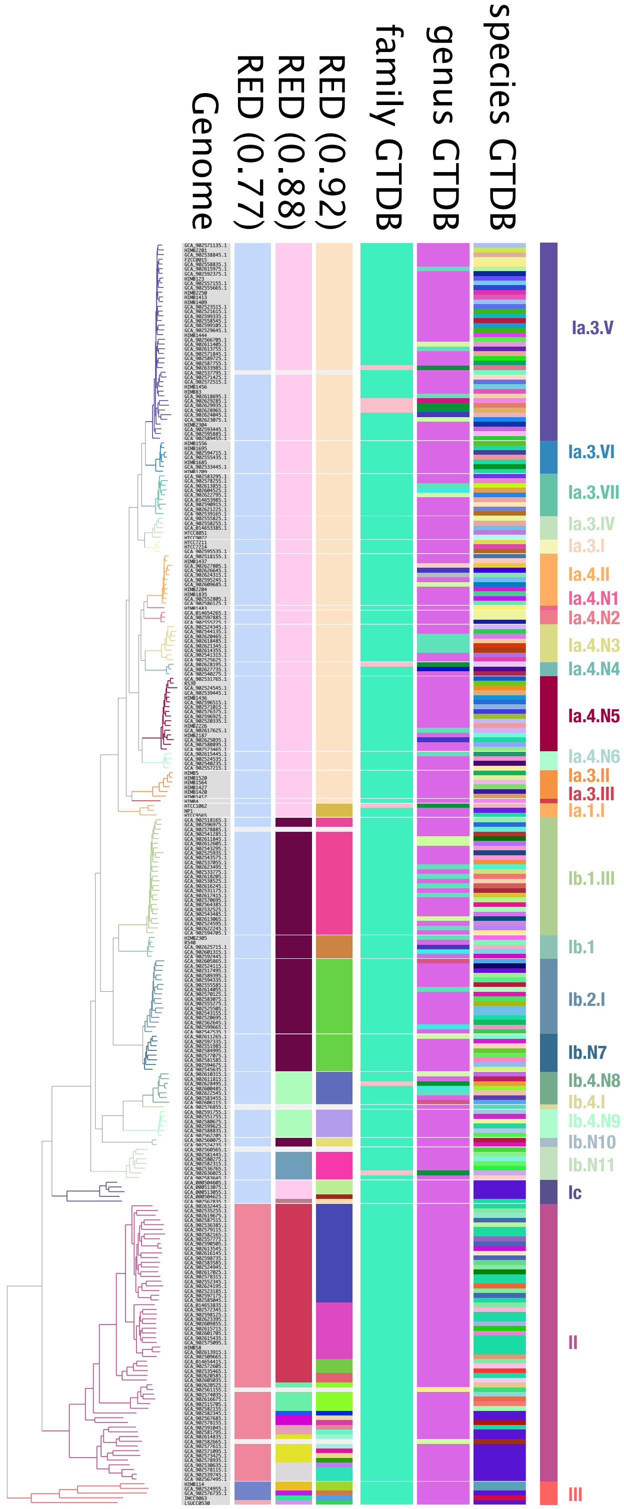


**Supplementary Fig. 9. Phylogenomic tree of the *Pelagibacterales* compared to other classification methods.** The phylogeny includes 268 genomes (50 isolates and 218 SAGs) included in the pruned phylogeny. Circles indicate ultrafast bootstrap support values ≥95% or greater from 1000 replicates.

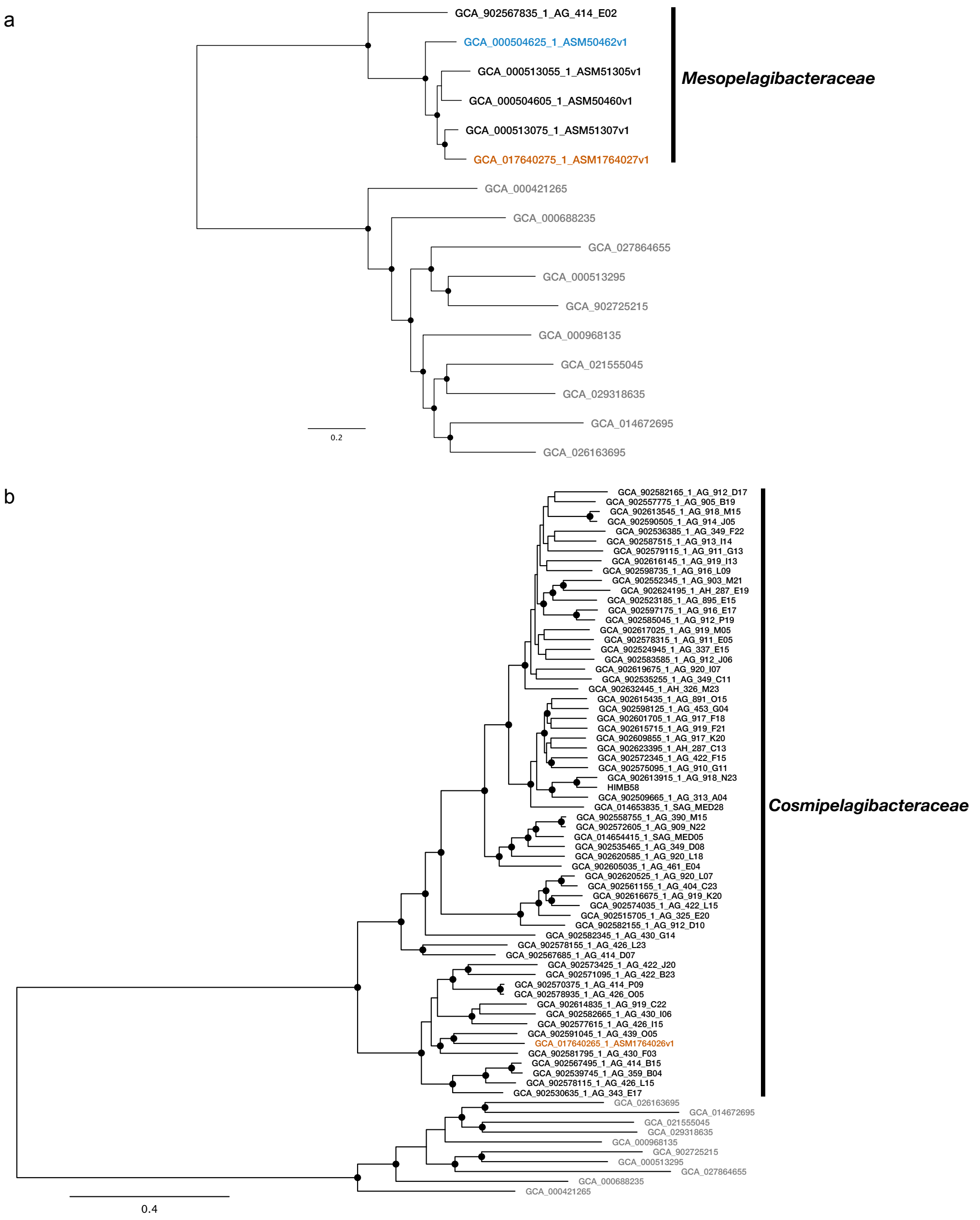


**Supplementary Fig. 10. Phylogenomic placement of previously described type material for 'Mesopelagibacter carboxydoxydans', *Mesopelagibacter profundi*, and *Anoxypelagibacter denitrificans*.** Phylogenies based on the SAR11_165 gene set for the (**A**) *Mesopelagibacteraceae* and (**B**) *Cosmipelagibacteraceae.* Both phylogenies include the single amplified genomes used for analyses in this study (black). The type genomes for 'Mesopelagibacter carboxydoxydans' (MAG GCA_017640275.1) and *Anoxypelagibacter denitrificans* (MAG GCA_017640265.1) are highlighted in orange, while *Mesopelagibater profundi* (SAG GCA_000504625.1) is highlighted in blue. The outgroup is shown in gray. Ultrafast bootstrap support values from 95 to 100 are indicated by circles at the relevant node.
